# A nucleic acid prodrug that activates mitochondrial respiration, promotes stress resilience, and prolongs lifespan

**DOI:** 10.1101/2024.11.16.623599

**Authors:** Takahisa Anada, Michiharu Kawahara, Taisei Shimada, Ryotaro Kuroda, Hidenori Okamura, Daiki Setoyama, Fumi Nagatsugi, Yuya Kunisaki, Eriko Kage-Nakadai, Shingo Kobayashi, Masaru Tanaka

## Abstract

Mitochondrial dysfunction caused by aging leads to decreased energy metabolism, resulting in functional decline and increased frailty in multiple tissues. Strategies for protecting and activating mitochondria under stressful conditions are required to suppress aging and age-related diseases. However, it is challenging to develop drugs capable of boosting mitochondrial respiration and compensating for the reduced intracellular adenosine triphosphate (ATP) levels. In this study, we developed a prodrug that stimulates the metabolism of intracellular adenine nucleotides (AXP: adenosine monophosphate (AMP), adenosine diphosphate (ADP), and ATP). It enhances AMP-activated protein kinase activity, fatty acid oxidation, oxidative stress resistance, and mitochondrial respiration, thereby increasing the intracellular ATP levels. Furthermore, this prodrug markedly extended the lifespan of *Caenorhabditis elegans*. AXP-driven stimulation of cellular energy metabolism proposed herein represents a novel geroprotective strategy and paves the way for the development of bioenergetic-molecule therapeutics.

## 1. Introduction

Aging and age-related diseases have been shown to be strongly associated with an imbalance between energy demand and supply^1^. Interventions such as calorie restriction, exercise, and molecules targeting longevity-related genes have been reported to be effective in addressing these issues^2^. Functions of mitochondria, the organelles responsible for energy production, decline with age, leading to age-dependent reductions in intracellular adenosine triphosphate (ATP) levels. There are many reports that ATP levels decrease with age in human and various animals, such as in human fibroblasts^3^, erythrocytes^4^, skeletal muscles^5^, rat heart tissues^6^, mouse retinas and brains^7^, *Drosophila*^8^, and *Caenorhabditis elegans*^9^ (*C. elegans*). ATP functions not only as an energy currency in biological systems, but also has diverse roles, including serving as a building block for ribonucleic acids, a donor in protein phosphorylation, and a solubilizing enhancer for maintaining proteostasis^10^. Therefore, its depletion might affect the maintenance of cellular functions. Mitochondrial dysfunction and decreased ATP levels impair energy metabolism and induce vulnerability in the entire body at the tissue level. Mitochondria play a crucial role not only in energy supply but also in the development of age-related diseases such as neurodegenerative and metabolic disorders^11–16^. Hence, there is a pressing need to develop techniques that protect against stress and activate the mitochondria. However, few drugs are capable of activating mitochondria and restoring reduced intracellular ATP levels, making the development of such mitochondria-activating drugs challenging. For this reason, this study hypothesized that by activating energy metabolism, particularly mitochondrial function, and increasing intracellular ATP concentration using pharmacological agents, it may be possible to slow the aging process.

ATP and its derivatives have been used as therapeutic agents to improve cardiovascular health, muscular performance, body composition, and fatigue recovery^17^. Despite the evidence of the therapeutic effects of ATP derivatives, their efficacy remains controversial. Some reports suggest that ATP administration is ineffective as a pharmaceutical agent^18^ because of the limited blood stability^19^ of adenosine derivatives and the lack of cell membrane permeability of the negatively charged phosphate groups, hindering entry into cells. Furthermore, cells recognize extracellular ATP as an inflammatory signal^20^. To minimize side effects, ATP administration techniques include drug delivery methods that encapsulate ATP in microparticles such as liposomes, micelles, and nanospheres^21^. Nevertheless, these approaches have limited utilization and applicability in the treatment of age-related diseases. Owing to the difficulty of increasing intracellular ATP concentrations, the long-term effects of ATP administration as a drug remain unclear.

Therefore, the present study was designed to develop the world’s first prodrug with enhanced biological stability and cellular permeability of adenosine phosphate, capable of stimulating AXP (adenosine monophosphate (AMP), adenosine diphosphate (ADP), and ATP)-associated metabolic pathways, boosting mitochondrial respiration and ATP production, thereby reducing aging.

The strategy of the present study involved developing an AXP prodrug (proAX, with its chemical structure shown in Figure 1a and synthetic scheme in Figure S1) that can be converted to ATP via AMP and ADP through intracellular enzymatic reactions and hydrolysis by phosphoramidating the phosphate moiety. Several phosphoramidate prodrugs (ProTides)^22^ have been successfully applied to mononucleoside analogs as a class of nucleoside-based antivirals, such as Sofosbuvir and Remdesivir, targeting hepatitis C virus^23^, and COVID-19^24^. Remdesivir is deemed safe for the treatment of hospitalized COVID-19 patients and approved by the Food and Drug Administration (FDA). These aryloxy phosphoramidate prodrugs are designed with the following proposed cellular activation pathway in the cell^25^: the prodrugs penetrate cell membranes, and the phosphoramidate moiety is metabolized stepwise by enzymatic and hydrolytic reactions to produce monophosphorylated nucleosides. These compounds are triphosphorylated by intracellular kinases and exert active medicinal effects. The potential intracellular activation mechanism of proAX developed in this study and based on this pathway is illustrated in Figure 1b. Moreover, we expected that, in addition to the direct conversion pathway of proAX to ATP, the transient increase in intracellular AMP levels might activate adenosine monophosphate-activated protein kinase (AMPK), which is involved in mitochondrial homeostasis regulation^26^. AMPK activation suppresses anabolic processes that consume ATP and promotes ATP synthesis through catabolic reactions.

**Figure 1.**
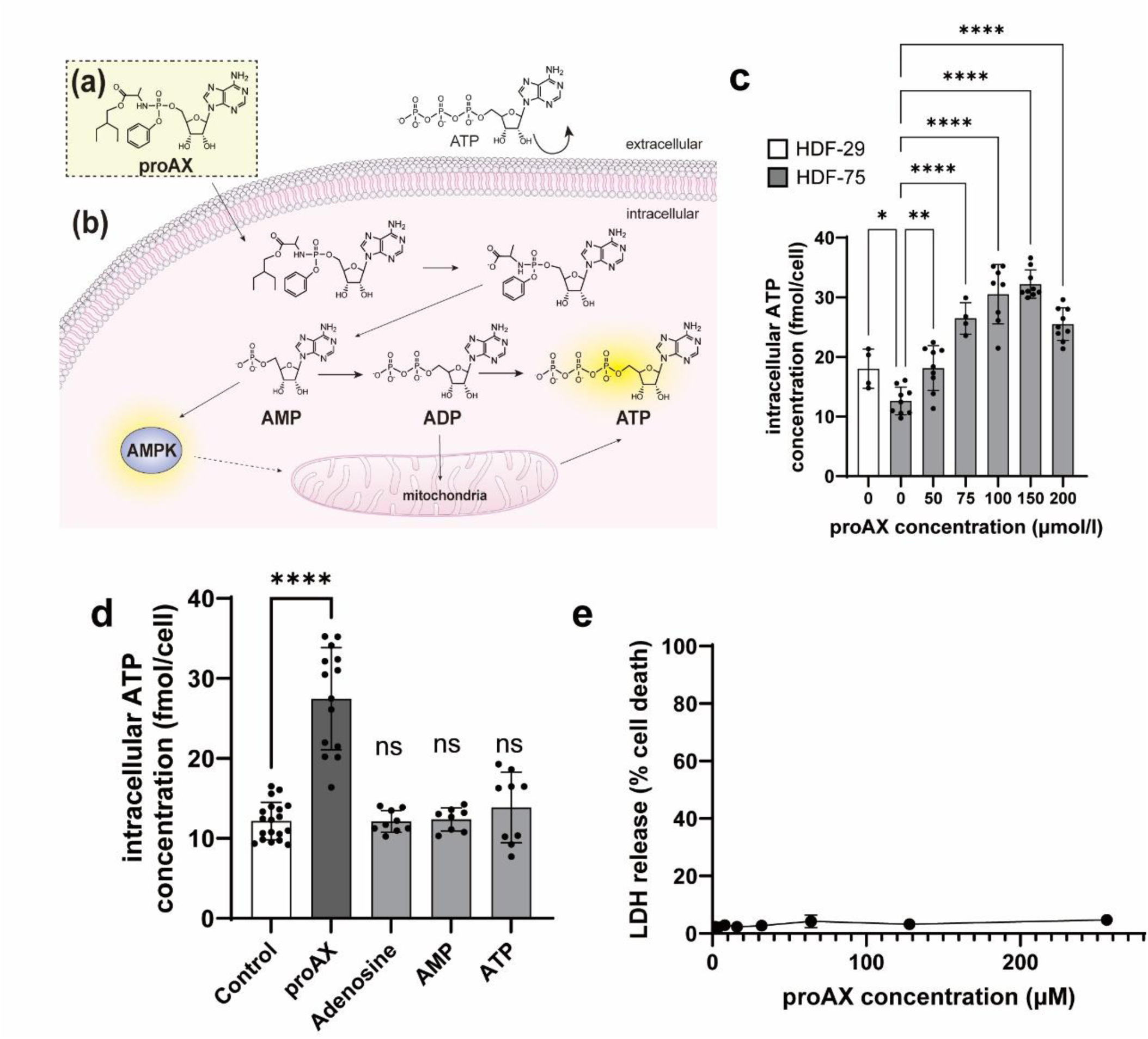
The strategy of this study, and proAX enhances intracellular ATP levels without inducing cytotoxicity. a, The chemical structure of proAX. b, ProAX is internalized into a cell and undergoes cellular metabolic reactions. ProAX is metabolized by intracellular enzymes, such as esterases and phosphoamidases, and is then converted into AMP, ADP, and ATP. Transient increases in AMP levels could stimulate AMP-activated protein kinase (AMPK). The activation of AMPK could promote the ATP synthesis pathways through catabolic pathways and improved mitochondrial function. c, Comparison of intracellular ATP concentrations between HDF-29 and HDF-75, and the effects of varying proAX concentrations on intracellular ATP levels in HDF-75. d, Changes in intracellular ATP concentrations upon addition of adenosine and its derivatives (100 µM each) to HDF-75. e, Impacts of proAX addition on HDF-75 as determined by a LDH assay. For panels c and d, mean ± standard deviation shown, along with statistical significances for relevant comparisons. Significance is assessed using a one-way ANOVA with Dunnett’s multiple comparison test, with *n* = 4–19 replicates per condition. * *p* < 0.05, \*\**p* < 0.01, \*\*\*\**p* < 0.0001; ns indicates not significant.

Research on ProTide technology has primarily focused on its conversion into synthetic compounds that do not exist in nature. These compounds exhibited antiviral and anticancer properties. However, to the best of our knowledge, no instances of ProTide-type drugs exhibiting activity in a form that perfectly matches the structure of natural nucleic acids have been reported. To our knowledge, this is the first report of a prodrug that enables intracellular conversion to AXP. In this study, we propose an innovative strategy for the “bioenergetic-molecule therapeutics” to activate mitochondrial respiration and increase intracellular ATP levels, which decrease with aging, disease, or both. We also demonstrated that proAX effectively extends the lifespan of *C. elegans*. We believe our findings open up new possibilities for nucleic acid prodrugs that achieve anti-aging through “stimulation of AXP-associated metabolic pathways (AXP stimulation)” distinct from existing anti-aging drugs.

## 2. Results

### 2.1. Capability of ATP augmentation and cytotoxicity of the prodrug

The results of culturing normal human dermal fibroblasts from 29-year-old and 75-year-old donors (HDF-29 and HDF-75) are displayed in Figure 1c. In the absence of the prodrug, HDF-75 cells exhibited lower ATP levels than HDF-29 cells, a phenomenon previously reported in the literature^3^. We examined whether proAX increased the intracellular ATP levels in HDF-75 cells. After incubation with proAX for 24 h, the addition of the synthesized proAX to HDF-75 cells led to a concentration-dependent increase in intracellular ATP levels. With 100 µM proAX, a 2.5-fold increase in ATP levels was observed compared to the control (absence of proAX). We found that proAX effectively elevated the ATP levels in the cells of elderly individuals with reduced ATP levels. Similar results were obtained in HDF-29 cells (Figure S3a). These results demonstrate that the prodrug can increase intracellular ATP levels regardless of donor age. Addition of ATP analogs or proAX to compare intracellular ATP levels showed that adenosine, AMP, and ATP did not change the intracellular ATP concentration; however, addition of proAX led to a significant increase (Figure 1d). These findings confirm the effectiveness of the newly developed proAX in increasing intracellular ATP concentrations, unlike conventional ATP analogs, which are less effective because of their limited stability and cellular membrane permeability. We assessed the cytotoxicity of proAX in HDF-75 cells by live/dead staining. The results showed that proAX exhibited very low cytotoxicity (Figure S3b). Furthermore, the lactate dehydrogenase (LDH) assay indicated that proAX was not cytotoxic up to 256 μM in HDF-75 cells (Figure 1e), although its addition appeared to suppress cell proliferation (Figure S3c, d).

### 2.2. Evaluation of time-dependent changes of AMP, ADP, and ATP concentration in the cells

We analyzed changes in the intracellular concentrations of AMP, ADP, and ATP after proAX administration to HDF-75 cells using liquid chromatography mass spectrometry (LC/MS). AMP and ADP levels increased significantly within 3 h of proAX administration and continued to increase over 72 h (Figure 2a, b). After 3 h of prodrug administration, AMP and ADP levels in the cells were approximately 3-fold and 1.5-fold higher, respectively, than those observed without the prodrug. Conversely, an increase in ATP levels was observed later than that in AMP and ADP, starting after 6 h, and remained significantly higher than that in the control condition without drug administration after 72 h (Figure 2c). These data indicated that when proAX was added at the start of the culture and incubated for three days without additional supplements, an increase in intracellular ATP levels was observed up to 72 h. Using these values, we calculated the adenylate energy charges in the cells with and without proAX administration at each time point. The energy charge was defined as (ATP +1/2ADP)/(ATP + ADP + AMP)^27^. Up to 24 h after drug administration, the energy charge in the prodrug group tended to be lower than that in the non-treated group; however, in both conditions, the value remained around 0.9, within maintaining a range under physiological conditions (Figure 2d). Furthermore, we calculated changes in the AMP:ATP ratio induced by the addition of proAX. As illustrated in Figure 2e, proAX administration resulted in a 2.4-fold increase in the AMP:ATP ratio after 3 h. Over time, these ratios decreased. We found that proAX administration increased the adenine nucleotide pool (Figure 2f). These findings suggest that prodrug addition temporarily increased AMP, ADP, and ATP concentrations. Early after their addition, proAX enhanced the AMP:ATP ratio without deviating energy from the physiological norms. This indicates that proAX induces a pseudo-state of energy deficiency in the cells during the initial phase of addition.

**Figure 2.**
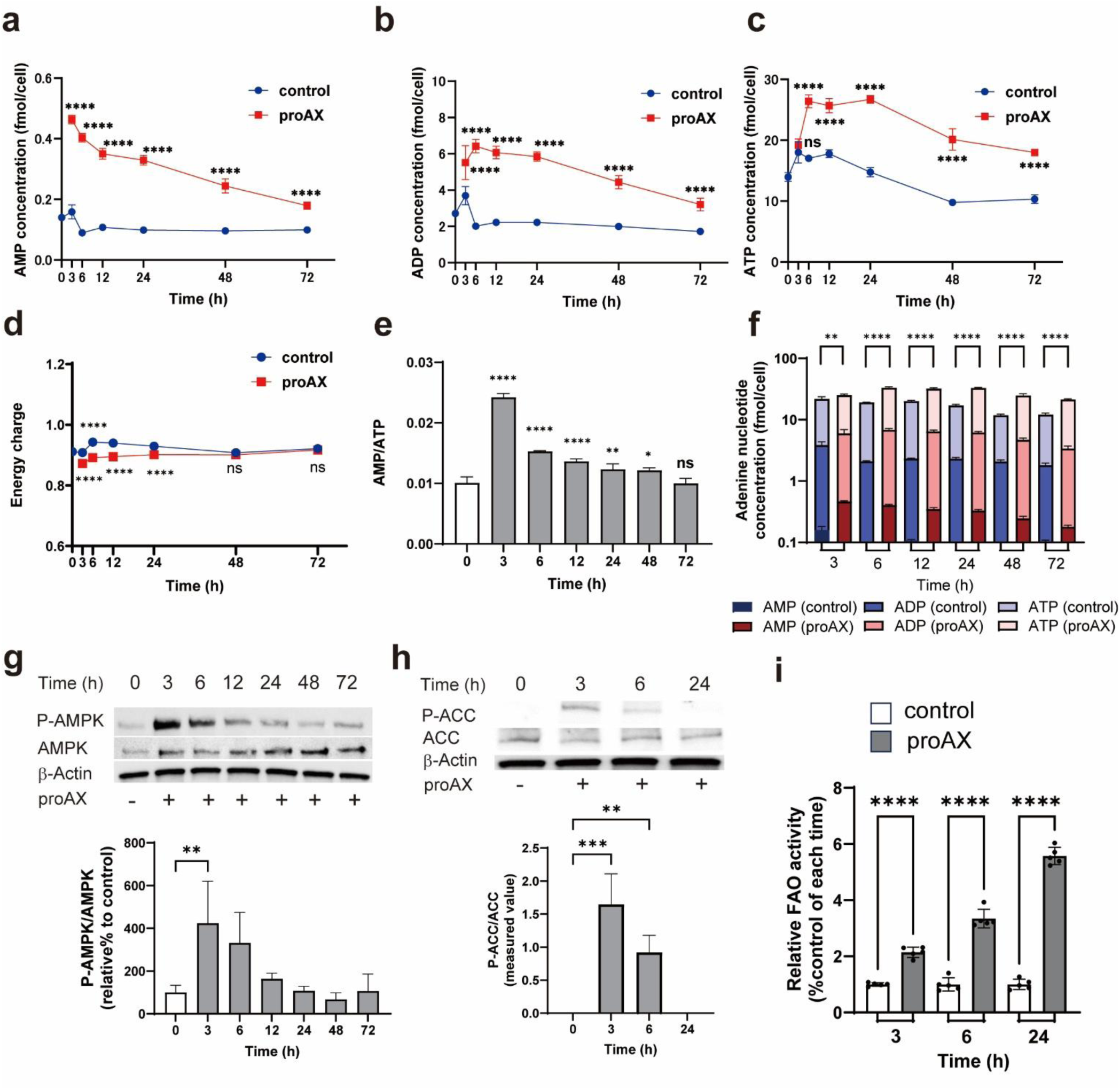
Time-dependent changes in intracellular AXP levels and reactions in HDF-75 following the addition of proAX. a, Changes in AMP concentration. b, Changes in ADP concentration. c, Changes in ATP concentration. d, Changes in adenylate energy charge. e, Changes in AMP:ATP ratio. f, Changes in adenine nucleotide concentration. g, Time-dependent changes in phosphorylation of AMPK induced by proAX in HDF-75. Representative western blots with anti-P-AMPK (Thr172) and anti-AMPK antibodies, h, Time-dependent changes in phosphorylation of ACC induced by proAX in HDF-75. Representative Western blots with anti-P-ACC, and anti-ACC antibodies, i, Relative fatty acid oxidation (FAO) activity in HDF-75 treated with proAX for 3, 6, and 24 h. For all panels, HDF-75 were treated with 100 µM proAX, mean ± standard deviation shown, along with statistical significances between relevant comparisons. Significance is calculated using a one-way ANOVA with Sidak’s multiple comparisons test (a-d, f). Panel e uses an ANOVA followed by Dunnett’s multiple comparisons test. \*\*\*\**p* < 0.0001, \*\**p* < 0.01, \**p* < 0.05, ns indicates not significant. For g, h, i panels, significance is calculated using a one-way ANOVA with Dunnett’s multiple comparisons test (g, h), Sidak’s multiple comparisons test (i). \*\*\*\**p* < 0.0001, \*\*\**p* < 0.001, \*\**p* < 0.01.

### 2.3. AMPK activation by proAX

The AMPK monitors cellular energy status. AMPK activation is triggered by an increase in the cellular AMP:ATP ratio. Given the changes in these ratios, as indicated in Figure 2e, suggesting AMPK activation, we investigated the activation of AMPK by proAX and its time dependency in HDF-75 cells. Immunoblot analysis revealed that AMPK phosphorylation occurred within 3 h of treatment with 100 μM proAX (Figure 2g). Phosphorylated AMPK levels increased approximately 4-fold 3 h after prodrug addition. AMPK activates fatty acid oxidation (FAO) by phosphorylating acetyl-CoA carboxylase (ACC), thereby reducing the levels of malonyl-CoA, an inhibitor of carnitine palmitoyltransferase (CPT). We assessed ACC phosphorylation after proAX administration (Figure 2h). ACC phosphorylation was observed within 3 h after the addition of proAX, and was almost undetectable within 24 h. Following the confirmed phosphorylation of AMPK and ACC, we further explored fatty acid catabolism, specifically FAO, using a fatty acid β-oxidation-responsive fluorescent probe. The results indicated that at 3, 6, and 24 h post-proAX administration, FAO activity was significantly increased compared to that in the control group (no drug added) (Figure 2i).

### 2.4. Intracellular mechanisms of the ATP boosting

First, we explored the intracellular uptake pathway of proAX. Changes in the intracellular ATP levels following the addition of proAX and the nucleoside transporter inhibitor dipyridamole are shown in Figure 3a. The observation that higher concentrations of the inhibitor partially inhibited the proAX-induced increase in intracellular ATP levels suggests that proAX is taken up by cells via both passive diffusion and nucleoside transporters.

**Figure 3.**
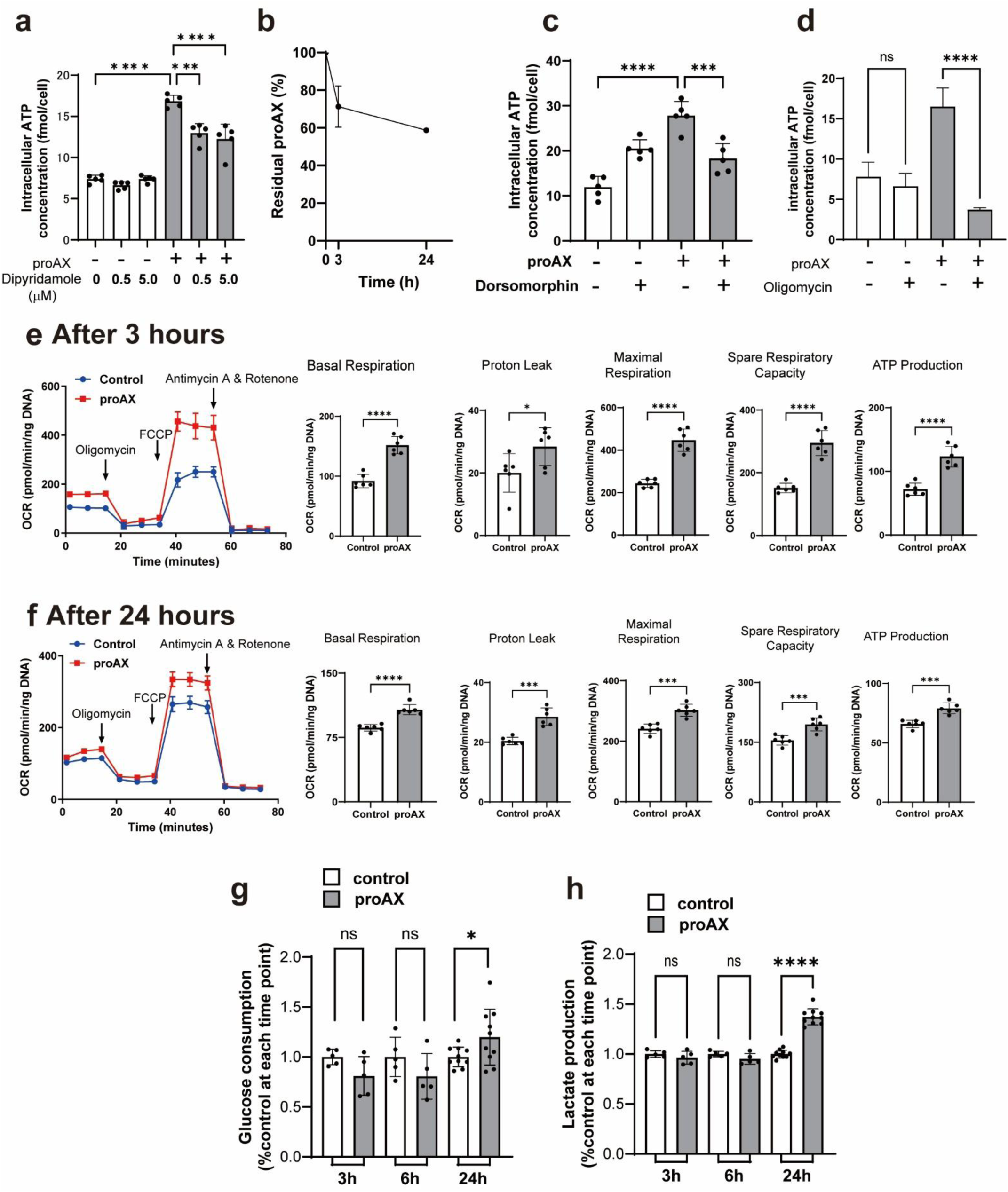
Mechanisms of augmentation of intracellular ATP by proAX administration. a, Intracellular ATP changes in HDF-75 treated with 100 µM proAX in the presence or absence of nucleoside transport inhibitor, dipyridamole, b, Quantification of residual [¹⁵N₂]-proAX in the culture medium after 3 and 24 hours of incubation. c, Intracellular ATP changes in HDF-75 treated with 100 µM proAX in the presence or absence of the AMPK-inhibitor compound-C (dorsomorphin), d, Intracellular ATP changes in HDF-75 treated with 100 µM proAX in the presence or absence of the ATP synthase inhibitor, oligomycin. The effects of proAX on mitochondrial respiration and their functions measured by an extracellular flux analyzer at 3 (e), and 24 (f) hours after treatment of 50 µM proAX. The effects of proAX on glucose consumption (g) and lactate production (h) at 3, 6, and 24 hours after treatment of 100 µM proAX. For all panels, mean ± sd shown, along with statistical significance between relevant comparisons. Significance is calculated using a one-way ANOVA with Dunnett’s multiple comparisons test (a-c). For panel d-g, significance was calculated using a two-tailed unpaired t-test. \*\*\*\**p* < 0.0001, \*\*\**p* < 0.001, \*\**p* < 0.01, \**p* < 0.05, ns indicates not significant.

To distinguish endogenous AXP from prodrug-derived AXP, we synthesized [¹⁵N₂]-proAX, in which two nitrogen atoms in the nucleobase moiety were labeled with the stable isotope ¹⁵N (Figure S2). The prodrug was added to HDF-75 cells, and the amount of proAX taken up by cells was estimated based on the residual proAX remaining in the culture medium after 3 and 24 hours (Figure 3b). Based on the amount of proAX remaining in the medium, we estimated that 28.7% of the total added proAX was estimated to be taken up by cells after 3 hours, and 41.3% after 24 hours. However, only about 1% of the intracellular proAX remained unmetabolized at 3 hours (Table 1). In a separate experiment, [¹⁵N₂]-proAX was added to HDF-75 cells, and intracellular levels of both unlabeled and [¹⁵N₂]-labeled AXP were measured. We found that the labeled AMP, ADP, and ATP accounted for approximately 48%, 35%, and 35% of their respective intracellular pools (Table 1). These results indicate that the proAX taken up by the cells was rapidly converted and metabolized within 3 hours.

**Table 1.** Intracellular AXP metabolite distribution after [¹⁵N₂]-proAX treatment.

|  | **[$^{15}\text{N}_2$]-proAX<sup>a</sup>** | **AMP** | **ADP** | **ATP** |
| --- | --- | --- | --- | --- |
| **[$^{15}\text{N}_2$]-labeled %** | < 1% | 47.7 ± 6.6% | 35.3 ± 0.9% | 35.0 ± 0.6% |

a. Percentage of intracellular prodrug remaining unmetabolized, relative to the total amount taken up by cells.

The increase in AMPK activity due to the addition of proAX was investigated to determine its effect on cellular ATP levels. This was evaluated by treating HDF-75 cells with dorsomorphin (Figure 3c), an AMPK inhibitor, and siRNA targeting AMPK (Figure S4a), followed by monitoring the changes in intracellular ATP concentration. ProAX-induced increase in cellular ATP levels was suppressed by dorsomorphin and siRNA. Furthermore, the phosphorylation of ACC by western blotting demonstrated that the treatment of proAX activated FAO processes. Collectively, these findings suggested that proAX activates AMPK, which in turn stimulates FAO, leading to increased intracellular ATP concentrations. For comparison, the effect of 5-aminoimidazole-4-carboxamide-1-D-ribofuranoside (AICAR), an AMPK activator, on intracellular ATP levels was examined in HDF-75 cells. However, no increase in the ATP concentration was observed following AICAR treatment (Figure S4c). These results suggest that AMPK activity is necessary for the proAX-induced increase in ATP levels, and that AMPK activity alone is insufficient.

To investigate how proAX increases intracellular ATP levels, changes in ATP levels were examined in HDF-75 cells after the addition of oligomycin, an inhibitor of ATP synthase (Figure 3d). In the absence of proAX, the addition of oligomycin did not significantly change the intracellular ATP levels. In contrast, with proAX, the addition of oligomycin led to a dramatic decrease in intracellular ATP levels.

Next, we investigated the effects of proAX on mitochondrial respiration using an extracellular flux analyzer. Figure 3e–f collectively demonstrate how proAX modulates mitochondrial respiration by using a Seahorse assay to measure the oxygen consumption rate (OCR) through sequential injections of agents such as oligomycin, FCCP, and rotenone/antimycin A, revealing that at an early time point (3 hours, as shown in Figure 3e), proAX alters basal respiration, ATP-linked respiration, and maximal respiration, and that these effects were maintained at 24 hours (as shown in Figure 3f). These findings indicate that proAX promotes a metabolic shift toward oxidative phosphorylation, thereby improving mitochondrial function and ATP generation, and elucidating its role in regulating cellular energy metabolism. Upon examining the changes in glucose consumption (Figure 3g) and lactate production (Figure 3h) before and after proAX administration, glucose consumption and lactate production were significantly increased 24 h after proAX administration. These results suggest that glycolytic metabolism was enhanced after 24 hours, implying that alongside improvements in oxidative phosphorylation (OXPHOS), the glycolytic system was also upregulated, which can be interpreted as proAX inducing a coordinated metabolic adaptation that enhances both energy pathways, thereby equipping the cells with overall ATP-generating flexibility to meet increased energetic demands.

### 2.5. The effects of proAX on oxidative stress resistance

Mitochondria generate cellular energy through OXPHOS; however, this process also produces reactive oxygen species (ROS) as byproducts. ROS damage the mitochondrial DNA (mtDNA), proteins, and lipids. To further investigate the impact of proAX on cellular robustness, its effects on oxidative stress resistance were examined in both cultured cells and *C. elegans*. Upon exposure to oxidative stress induced by hydrogen peroxide (Figure 4a), a comparison of oxidative stress resistance between young and aged cells revealed that at all hydrogen peroxide concentrations, HDF-75 consistently had lower cell counts than HDF-29. This indicates a reduced resilience to oxidative stress in aged cells. Pretreatment of HDF-75 cells with proAX before adding hydrogen peroxide mitigated the decline in cell number, yielding a reduction in cell viability compared to that of HDF-29 cells. This suggests that proAX treatment enhances the resilience of aged cells to oxidative stress.

**Figure 4.**
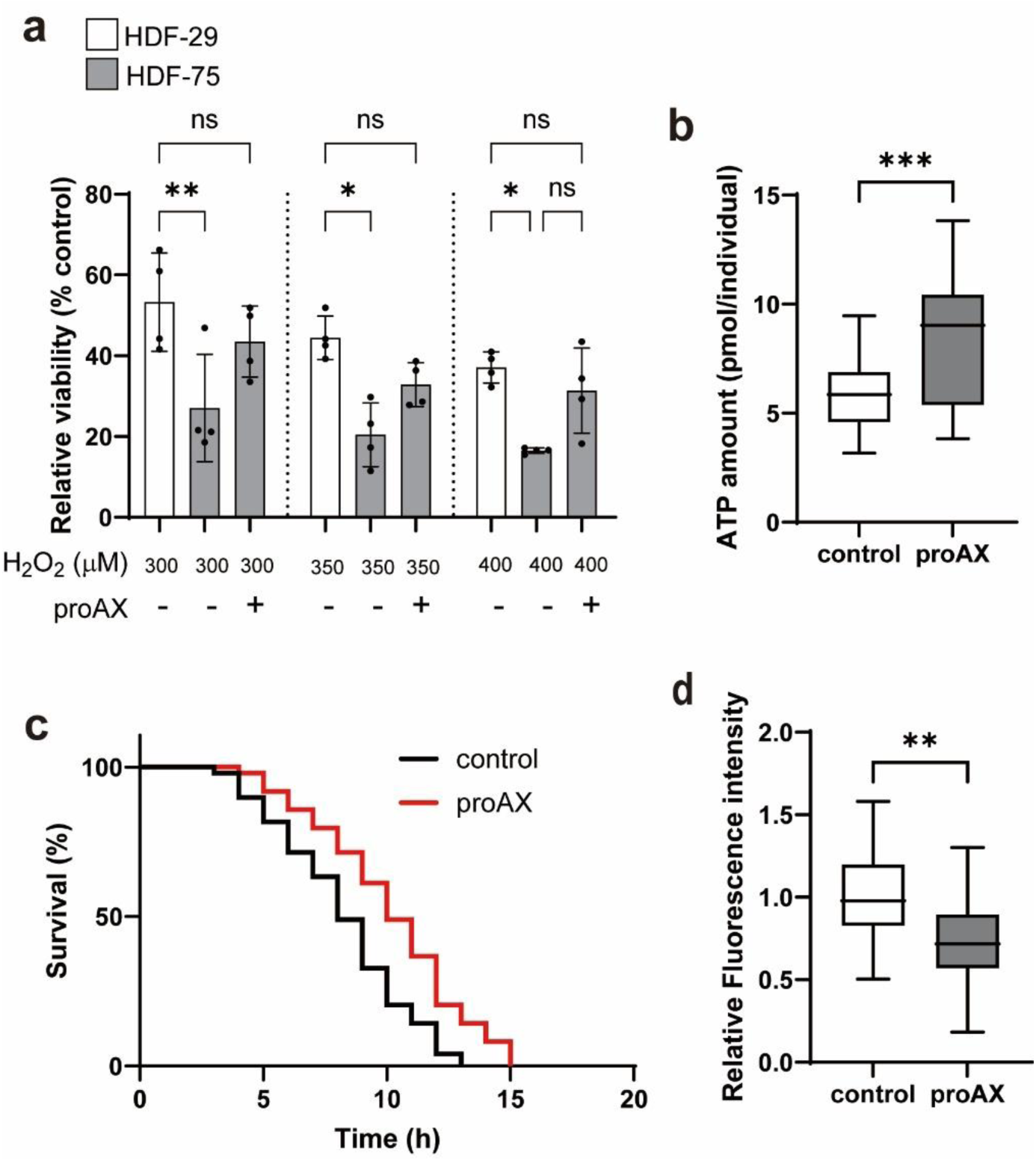
Effect of proAX on oxidative stress resistance in fibroblast cells and *Caenorhabditis elegans*. a, Fibroblasts derived from donors of different ages were treated with indicated concentrations of hydrogen peroxide following pretreatment with or without proAX (100 µM). b, ATP levels in *C. elegans* on Day 7 when cultured with or without proAX (100 µM) (n = 25-35 worms). c, Effect of paraquat-induced stress on *C. elegans* pretreated with proAX (100 µM) for 7 days (n = 49 worms, each). d, Relative ROS levels in *C. elegans* on Day 7 when cultured with or without proAX (100 µM) (n = 22-24 worms). For all panels, mean ± standard deviation shown, along with statistical significance between relevant comparisons. For the panel b and d, the center line of the box shows the median, box limits show interquartile ranges, and whiskers show the range of the data. Significance is calculated using one-way ANOVA with Tukey’s multiple comparisons test (a), a two-tailed unpaired t-test (b, d), and log-rank test (c). \*\*\**p* < 0.001, \*\**p* < 0.01, \**p* < 0.05, ns indicates not significant.

Paraquat, a mitochondrial toxicant, affects electron transport chain function through redox reactions, thereby increasing superoxide production^28^. To investigate whether proAX exhibits antioxidant stress effects at the organismal level, we selected *C. elegans* as the model organism. First, we examined whether proAX enhances the in vivo ATP levels at the organismal level in nematodes (Figure 4b). ProAX addition significantly increased the ATP levels per worm. These results clarify that proAX can increase ATP levels not only in cultured cells but also in individual organisms.

Here, the effects of proAX on oxidative stress resistance were examined by exposing 7-day-old *C. elegans* transferred to a NGM medium containing proAX to 200 mM paraquat. The results indicate that *C. elegans* cultured in the presence of proAX had a significantly extended lifespan (Figure 4c). Extension rate was 23%. Consequently, proAX enhances resilience to oxidative stress at the organismal level.

It has been reported that impaired energy metabolism leads to increased mitochondrial ROS production^29^, whereas improved mitochondrial function is associated with reduced ROS levels^30^. Based on these observations, we investigated the effect of the prodrug on endogenous ROS levels in *C. elegans*. As shown in Figure 4d, proAX treatment significantly reduced endogenous ROS levels in *C. elegans*.

A summary of proAX effects on cells is shown in summarized in Figure 5.

**Figure 5.**
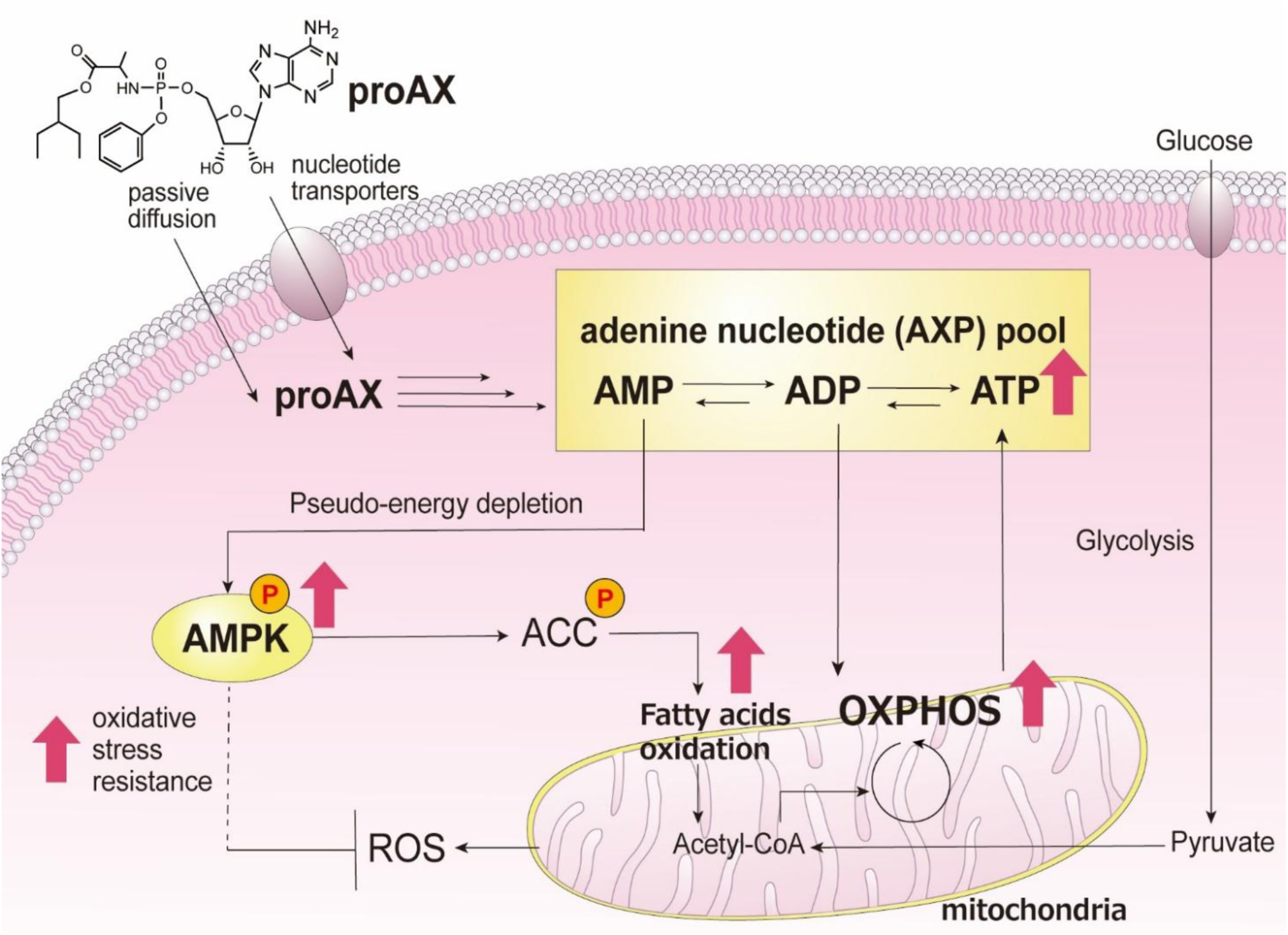
Summary of the effects of proAX on intracellular energy metabolism. ProAX penetrates cells, increases the AXP pool, and activates AMPK. The activation of AMPK promotes fatty acid oxidation and stimulates mitochondrial respiration. Furthermore, proAX is observed to enhance resistance to oxidative stress.

### 2.6. The effects of proAX on the lifespan of *C. elegans*

As *C. elegans* experiences a significant decrease in intracellular ATP levels with aging^9^, it could be a suitable model for confirming the effects of ATP elevation. *C. elegans* were treated with various concentrations of proAX, ranging from 25 to 300 μM, and their lifespans were measured (Figure 6a, and Figure S5a-d). Surprisingly, proAX concentrations over 50 μM resulted in significantly increased longevity compared to that in the control group without proAX. Figure 6a shows the survival curves of *C. elegans* treated with 100 μM proAX and without the prodrug (control). In contrast, AMP, ATP, or AICAR did not significantly extend the lifespan of *C. elegans* (Figure 6b). Table 2 summarizes and compares the effects of proAX, AMP, ATP, and AICAR on lifespan extension. When varying the concentration of proAX, 100 μM proAX was found to have the highest efficacy in extending lifespan (extension rate 24%, *p* < 0.0001). These results demonstrated the efficacy of proAX in extending the lifespan of *C. elegans*. To exclude the possibility of dietary restriction (DR) effects from proAX supplementation, worm body size was measured, as DR is known to increase longevity and reduce the growth of *C. elegans*. Furthermore, drugs that activate AMPK, such as metformin, have been reported to extend the lifespan of *C. elegans*, while inhibiting their growth^31^. There were no significant differences in body length between the treatments with or without proAX from Days 1 to 3 (Figure 6c). This was also the case for the body length differences between Days 4 and 20 (data not shown). Using dead bacterial lawns in lifespan experiments helps elucidate the impact of bacterial metabolism on the lifespan of *C. elegans*. Therefore, the bacteria were killed by heat, and lifespan analysis was performed in the same manner as for live bacteria. A significant lifespan extension effect of proAX was observed even when feeding on dead *Escherichia coli* (*E. coli*) (Figure S5e, and Table S1). This finding indicates that the effect of proAX is not dependent on *E. coli* or its metabolism.

**Figure 6.**
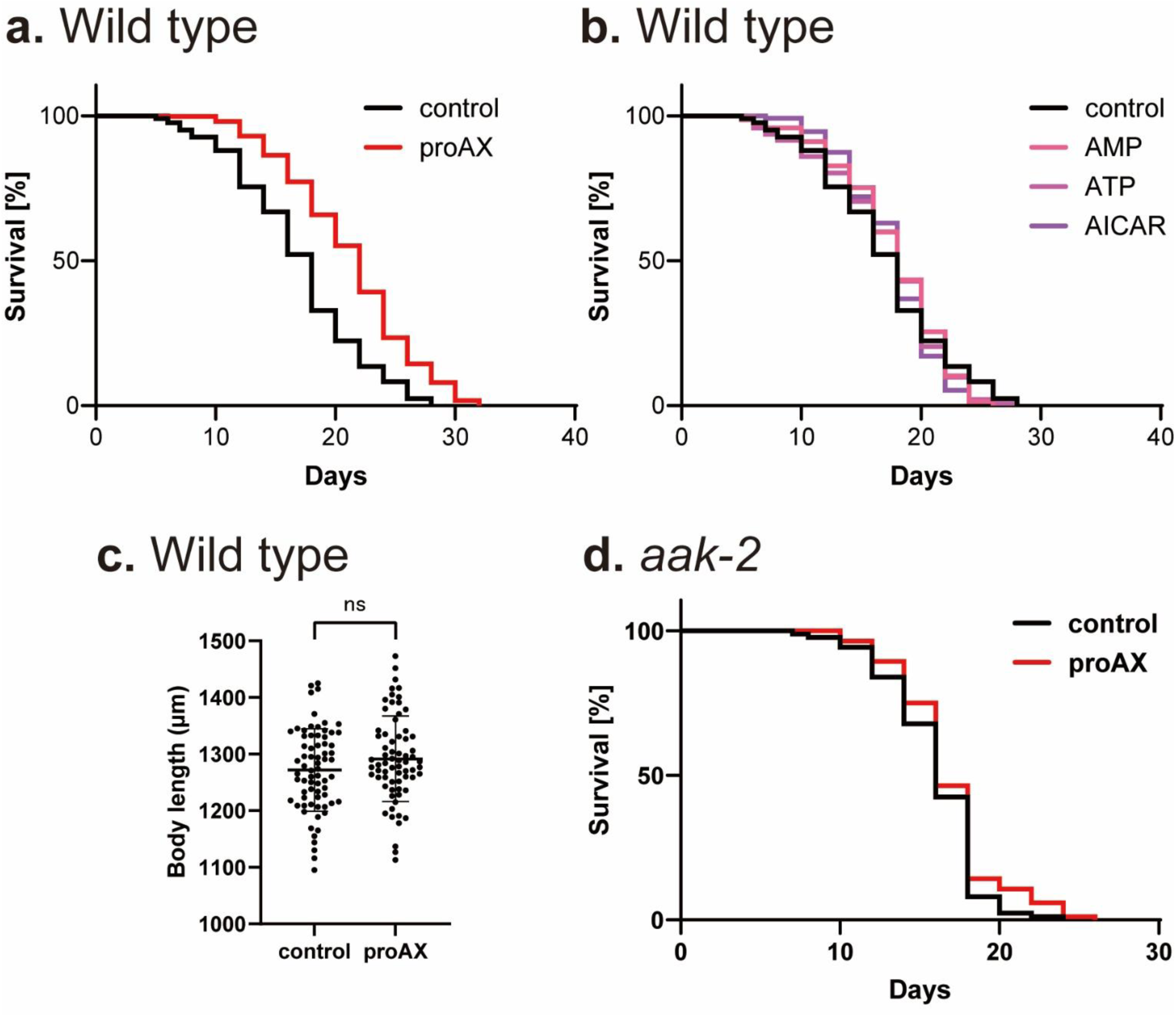
ProAX extends lifespan of *Caenorhabditis elegans*. a, Survival curves of *C. elegans* treated with proAX (100 µM). b, Survival curves of *C. elegans* treated with AMP, ATP, and AICAR (100 µM each). c, Effect of proAX on the growth of *C. elegans* over three days. d, Survival curves of *aak-2* mutant treated with proAX (100 µM). For all panels, mean ± standard deviation shown, along with statistical significance between relevant comparisons. Significance is calculated using log-rank test (a, b, d), a two-tailed unpaired t-test (c), and ns indicates not significant.

**Table 2.** Effect of proAX and other drugs on lifespan extension of *C. elegans*.

| sample | Mean lifespan ± SE (days) | Extension rate (%) | N | *p* value (log-rank) |
| --- | --- | --- | --- | --- |
| Control (no drug) | 17.5 ± 0.3 | - | 326 | - |
| 25 μM proAX | 18.1 ± 0.4 | 3.4 | 116 | 0.720 |
| 50 μM proAX | 19.4 ± 0.5 | 10.9 | 111 | 0.002 |
| 100 μM proAX | 21.7 ± 0.2 | 24 | 513 | $< 10^{-4}$ |
| 200 μM proAX | 19.3 ± 0.5 | 10.3 | 95 | 0.004 |
| 300 μM proAX | 20.2 ± 0.6 | 15.4 | 69 | 0.0002 |
| 100 μM AMP | 18.1 ± 0.4 | 3.4 | 145 | 0.823 |
| 100 μM ATP | 17.7 ± 0.4 | 1.1 | 142 | 0.891 |
| 100 μM AICAR | 18.0 ± 0.4 | 2.8 | 111 | 0.715 |

To further investigate the mechanism underlying the lifespan-extending effect of proAX, we examined its effect on an *aak-2* loss-of-function mutant, which lacks the *C. elegans* ortholog of AMPK (Figure 6d). As a result, proAX did not extend the lifespan of the *aak-2* mutant (Table S2). These findings suggest that the AMPK signaling pathway plays an important role in mediating the proAX-induced extension of lifespan.

## 3. Discussion

The relationship between aging and energy metabolism has long been the subject of discussion. Among various factors that contribute to aging, mitochondrial dysfunction plays a pivotal role in the aging process. This dysfunction is also associated with the onset of age-related diseases. Small molecules essential for life activities such as NAD^+^ and taurine^32,33^, have been shown to slow aging progression in various animals. Reports suggest that supplementing these molecules from outside the body can extend the lifespan. Similarly, ATP levels decrease with age. The decline in ATP levels associated with reduced energy metabolism is thought to disrupt energy homeostasis and potentially accelerate aging. However, there are few reports on the effective supplementation of intracellular ATP levels with drugs. Various genetic mutations are known to extend the lifespan of *C. elegans*, with a significant increase in ATP levels observed in many mutant strains^14^. Kipreos et al. suggest that while genetic mutations increasing ATP levels benefit longevity, they propose the “energy maintenance theory of aging (EMTA)” as not extending longevity but as essential for survival maintenance in older animals^14^. Our results illuminate how AXP energy metabolism stimulation could provide a new paradigm for the role of energy metabolism in aging, particularly through its involvement in mitochondrial activity, the AXP pool, and the energy sensor AMPK.

We demonstrated that synthesized proAX increased intracellular ATP levels (Figure 1c). We found that the number of cells, 24 h after proAX administration, decreased in a concentration-dependent manner (Figure S3c, d). LDH and live/dead assays revealed no cytotoxicity of proAX (Figure 1e and Figure S3b), suggesting that the decrease in cell number was due to the suppression of cell proliferation. Numerous studies have shown that AMPK activators, such as metformin and AICAR, inhibit cell proliferation through AMPK activation^34,35^. AMPK conserves energy and enhances ATP utilization efficiency by inhibiting energy-intensive processes such as protein synthesis and cell proliferation. We confirmed that AICAR inhibited the proliferation of HDF-75 cells (Figure S4b). AMPK activation promotes catabolic pathways to restore energy homeostasis and enhance ATP production. However, addition of the conventional AMPK activator AICAR to HDF-75 cells did not increase intracellular ATP levels within 24 h (Figure S4c).

As expected from the molecular design, proAX was first converted into AMP in the cell, which significantly increased its concentration. Subsequently, the cellular levels of ADP and ATP increased (Figure 2a-c). Consequently, the intracellular reactions triggered by proAX increased the adenine nucleotide pool (Figure 2f). The energy charge stayed around 0.9 from 3 to 72-h post-proAX administration, indicating no abnormal cellular energy deficiency or excess (Figure 2d). AMPK activation is generally triggered by decreased energy status or stress. However, energy-charge fluctuations during proAX administration suggested that the cells did not experience significant energy stress, implying that a temporary increase in AMP concentration (AMP/ATP) caused by proAX induced AMPK activation (Figure 2g). The elevation of AMP and ADP levels by proAX may protect AMPK from dephosphorylation, potentially sustaining its activity longer^36^. This indicates that the prodrug may cause the cells to perceive a transient pseudo-energy depletion.

The intracellular uptake pathway and the activation mechanism of the prodrug were examined. ProTide technology was developed to efficiently deliver nucleotide analogs by enhancing cell membrane permeability^37^. Investigations into the contribution of nucleoside transporters to the intracellular uptake of proAX suggested that, in addition to passive diffusion, some proportion of the prodrug was internalized via the transporters (Figure 3a), along with other nucleoside drugs for cancer chemotherapy^24^. ProAX, once internalized, undergoes metabolic conversion similar to those of other ProTides. It is transformed into an alanine intermediate through esterase hydrolysis and converted into its monophosphate form (AMP) by phosphoamidases.

Using the stable isotope-labeled prodrug, we estimated the proportion of intracellular AXPs derived from the proAX (Table 1). AXPs are rapidly and continuously interconverted and metabolized within short timeframes. ATP participates in a wide variety of cellular processes, including the synthesis of adenine derivatives and other purine bases, as well as incorporation into coenzyme A, NAD⁺ derivatives, post-translational modifications, and macromolecules such as mRNA. Due to the broad involvement of ATP in these pathways, it was difficult to comprehensively trace all downstream metabolites of the prodrug. However, the observed uptake of [¹⁵N₂]-proAX into cells, along with a measurable increase in isotope-labeled AXPs, provides evidence that the prodrug is converted and metabolized within 3 hours. These results support the conclusion that the prodrug contributes to the elevation of intracellular AXP levels either directly or via its metabolic intermediates.

AMPK activation inhibits anabolic pathways and reduces ATP consumption. Simultaneously, it activates catabolic pathways that enhance ATP production via β-oxidation and glycolysis. Studies have shown that AMPK activation in the heart^38^ and monocytes^39^ leads to activation of the glycolytic pathway. It has been reported that AMPK increases the flux of the glycolytic pathway^40^. In this study, glucose consumption and lactate production did not increase during short culture periods (3, and 6 h), but both increased after 24 h (Figure 3g, h). This suggests that the non-oxygen-dependent glycolytic pathway may be promoted by a sufficient increase in ATP levels. Kobayashi et al. demonstrated that AMPK activation increased fatty acid oxidation, leading to elevated mitochondrial respiration^41^. Our results suggest that AMPK activation enhances FAO (Figure 2i), thereby improving ATP production. However, as ATP production was not enhanced by AMPK activators such as AICAR (Figure S4c), it is conceivable that proAX not only activates AMPK, but also enhances ATP production through other mechanisms.

Several enzymes, including adenylate kinases, phosphorylate antiviral nucleotide analogs and natural AMP to ADP. Subsequently, these diphosphorylated drugs are further phosphorylated by multiple enzymes, including nucleoside diphosphate kinases, phosphoglycerate kinases, pyruvate kinases, and creatine kinase^42^. In the case of proAX, after being metabolized by AMP, these enzymes can convert AMP to ADP, and ADP to ATP. Although it is not yet possible to estimate the proportions of these reactions that contribute to ATP production accurately, experiments using oligomycin, an ATP synthase inhibitor, have shown that ATP synthesis is significantly suppressed (Figure 3d). This suggests that ATP production by proAX in the mitochondria is predominant. A close relationship exists between adenosine homeostasis, adenine nucleotide pool size, and mitochondrial bioenergetics^43–45^. When the concentration of ADP in the cell increases, it is transported into the matrix through the ADP/ATP exchange transport system, promoting ATP synthesis. ADP regulation of respiration leads to increased oxygen consumption in the mitochondria^46^. In other words, the prodrug likely boosted ADP levels, serving as a substrate for oxidative phosphorylation, thereby enhancing ATP production via mitochondrial respiration (Figure 3e, f). Kamatani et al. suggested that enhancing the purine salvage pathway, thereby reducing de novo purine synthesis, decreasing energy expenditure, and increasing the purine pool, leads to enhanced cellular ATP^47^. The critical role of mitochondria in cellular processes is well-established, with evidence linking mitochondrial dysfunction to aging and age-related diseases^48,49^. Strategies to enhance mitochondrial quality and function show the potential for widespread benefits, including aging and age-related disease prevention and treatment, as well as for improving overall cellular health and longevity. Our results demonstrate that the administration of proAX significantly enhances the energy metabolism of HDF cells via OXPHOS. This enhancement of energy metabolism may result from enhanced fatty acid oxidation due to AMPK activation and an increased adenine nucleotide pool available for ATP synthase.

Studies have shown that the intracellular adenine nucleotide pool correlates with the longevity of organisms such as yeast^50^, *C. elegans*^51,52^, and *Drosophila*^53^. In these organisms, lifespan extension is often linked to genetic mutations that increase the AMP/ATP ratio by affecting the adenine nucleotide pool, thereby activating AMPK.

Therapeutic approaches that activate AMPK, such as caloric restriction and metformin, have been shown to improve longevity and healthspan^54,55^. The effects of AMPK are elicited through mechanisms involving interactions with various factors. However, other studies have reported that AICAR does not extend *C. elegans*’s lifespan^56^. Our experiments confirmed that AICAR administration did not extend the lifespan of *C. elegans* (Figure 6b). Therefore, although AMPK activation influences the lifespan extension in *C. elegans*, its effect alone may be insufficient, indicating the involvement of other factors.

Compared to existing AMPK activators such as metformin and AICAR, proAX is distinguished by its dual functionality in AMPK activation and mitochondrial enhancement, as well as by its hydrophobic character. This compound transiently elevates intracellular AMP levels, which activates AMPK and dynamically engages the cellular energy-sensing pathway. Additionally, the concomitant increase in ADP levels, which serve as substrates for ATP synthesis, is expected to rapidly stimulate ATP-producing pathways. Notably, proAX exerts a more pronounced effect on intracellular AMP and ADP concentrations than either metformin or AICAR.

Unlike metformin, which indirectly activates AMPK by inducing a transient energy-deficient state, proAX appears to activate the AMPK pathway in the absence of energy stress. This property may offer a distinct advantage, as it enables enhancement of ATP levels and overall energy status, potentially supporting or even improving cellular function while selectively promoting beneficial AMPK-mediated signaling. The dual modulation of AMPK signaling, and mitochondrial function may underlie the enhanced physiological outcomes observed with proAX treatment.

Our findings raise the question: What impact does enhancing intracellular ATP levels have on the lifespan of *C. elegans*? Insulin receptor mutants in *C. elegans* have been shown to promote mitochondrial fusion, increase ATP levels, and extend the lifespan^57^. Although few methods are available to pharmacologically enhance ATP levels, some studies have explored the relationship between increased ATP levels and the lifespan of *C. elegans*. Higashitani et al. reported that MA-5, which binds to ATP synthase and enhances ATP production, does not extend *C. elegans*’s lifespan^58^. Other reports suggest that a combination of febuxostat and vitamin C, an antioxidant, favors the purine salvage pathway, thereby aiding ATP production and extending the lifespan of *C. elegans* through antioxidant effects, which can be considered supportive of our study’s findings^59^. In the present study, we demonstrated that proAX, without the addition of antioxidants, such as vitamin C, independently enhanced antioxidant resistance in both fibroblasts and *C. elegans* (Figure 4a, c).

Furthermore, the observed reduction in endogenous ROS levels in *C. elegans* following proAX treatment suggests that the prodrug may enhance the efficiency of the mitochondrial respiratory chain, leading to increased ATP production and suppressed ROS generation. In addition to mitochondrial function and elevated ATP levels, multiple mechanisms have been reported to contribute to enhanced oxidative stress resistance, including regulation of apoptosis, ATP-dependent molecular chaperones, and antioxidant enzyme systems. With respect to apoptosis, sufficient intracellular ATP levels favor energy-dependent apoptotic pathways, enabling the controlled removal of damaged cell. In contrast, ATP depletion can impair apoptotic signaling and promote necrosis, a process often accompanied by the release of inflammatory cytokines and increased oxidative stress^3,60^. Another potential mechanism involves ATP-dependent molecular chaperones, which are upregulated in response to external stress and function to preserve cellular homeostasis. For instance, HSP70, a well-characterized chaperone, requires ATP to mediate protein folding and protect cellular integrity^61^. Moreover, activation of the AMPK–FOXO3 pathway has been reported to induce the thioredoxin system, a major antioxidant defense mechanism implicated in the reduction of ROS levels^62^. It is therefore plausible that proAX may influence this pathway. In addition, AMPK activity and overall energy metabolism have been linked with the expression of Nrf2, a transcription factor that regulates genes encoding antioxidant enzymes in response to oxidative stress^63^. However, it should be noted that the efficacy of antioxidant supplementation in extending lifespan remains controversial^64^. Therefore, further studies are warranted to clarify the complex relationship between oxidative stress and aging, as well as to elucidate the precise mechanisms through which proAX exerts its effects.

## 4. Conclusion

Overall, proAX, which is converted into a natural energy currency, increases the intracellular adenine nucleotide pool. It exhibits various effects, including AMPK activation, enhanced FAO, and activation of mitochondrial respiration, resulting in increased ATP production and improved resistance to oxidative stress. These combined actions are believed to significantly extend the average lifespan of *C. elegans*. Therefore, the novel concept proposed by this technology, AXP energy metabolism stimulation (AXP stimulation), may inhibit the decline in energy metabolism and the biological defense mechanisms associated with aging, thereby preventing the onset of age-related diseases. Although further studies are necessary to understand how proAX extends the lifespan of *C. elegans*, the results of this study support the significant role of the adenine energy metabolism pathway in controlling lifespan, suggesting that drugs targeting these pathways may influence lifespan.

## 5. Experimental Section

### Synthesis of proAX

ProAX and [^15^N_2_]-proAX were synthesized, and its chemical identity was established using nuclear magnetic resonance (NMR) and fast atom bombardment mass spectrometry (FABMS) analyses, as described in the Supplementary Materials.

### Cell cultures

Human dermal fibroblasts (HDFs) derived from 29- or 75-year-old donors were obtained from Cell Applications, Inc. (San Diego, CA, USA). Normal human dermal fibroblasts (NHDFs) were obtained from Lonza (Basel, Switzerland). HDF were grown at 37 °C and 5 % CO_2_ in HDF growth medium (Cell Applications, Inc., San Diego, CA, USA). NHDFs were grown at 37 °C and 5 % CO_2_ in DMEM/F12 supplemented with 10% FBS and 1% penicillin (all from Gibco, Carlsbad, CA, USA). ProAX was added at 3 h after seeding.

### Reagents

Chemicals used in this study: Adenosine 5’-monophosphate (#017-20391, Fuji Film Wako Pure Chemicals, Osaka, Japan), Adenosine 5’-triphosphate (#019-09672, Fuji Film Wako Pure Chemicals, Osaka, Japan), AICAR (#015-22531, Fuji Film Wako Pure Chemicals, Osaka, Japan), Dipyridamole (#D2274, TCI, Tokyo, Japan), Dorsomorphin (#D5349, TCI, Tokyo, Japan), Etomoxir sodium salt hydrate (#E1905, sigma, St. Louis, MO, USA), Hydrogen Peroxide (#H1222, TCI, Tokyo, Japan), Oligomycin (#G270, Dojindo, Kumamoto, Japan), Paraquat (#D3685, TCI, Tokyo, Japan).

### Antibodies

Antibodies used in this study: Anti-AMPKα (#2532; 1:1000), Anti-pThr172-AMPK (#2535; 1:1000), Anti-Acetyl-CoA Carboxylase (#3676; 1:1500), Anti-Phospho-CoA Carboxylase (#11818; 1:1500) from Cell Signaling Technology. Horseradish peroxidase-conjugated anti-rabbit IgG (#111-035-144; 1:10000) from Jackson ImmunoResearch.

### Intracellular ATP Levels assay by Luciferin-luciferase

Intracellular ATP levels in HDFs incubated with different concentrations of proAX were measured using a luciferin–luciferase-based ATP assay kit (Dojindo, Kumamoto, Japan) according to the manufacturer’s instructions. HDFs were seeded at 5,000 cells/well in 96-well plates. After 24 h of proAX treatment, cells were lysed in 0.2% Triton X-100 solution and sonicated in an ice bath. DNA concentration in cell lysate was measured using a Quant-iT PicoGreen dsDNA kit (Thermo Fisher Scientific, Waltham, MA, USA). The luciferin-luciferase luminescence was measured using an Infinite 200PRO M Plex microplate reader (Tecan, Zürich, Switzerland). The ATP levels was normalized using the cell number, as determined by the DNA concentration.

### Cytotoxicity Assay

The cytotoxicity of NHDF and HDF derived from a 75-year-old donor incubated with different concentrations of proAX was measured using a lactate dehydrogenase (LDH) assay kit (Dojindo, Kumamoto, Japan) or a Cellstain double staining kit (Dojindo, Kumamoto, Japan) according to the manufacturer’s instructions. For the LDH assay, NHDFs were seeded at 5,000 cells/well in 96-well plates. After 24 h of treatment with proAX, the absorbance of the cells was measured at 490 nm using an Infinite 200PRO M Plex microplate reader (Tecan, Zürich, Switzerland). For Live/Dead cell staining, HDF were seeded at a density of 20,000 cells/well into 96-well plates. After 3 h of treatment with proAX, images were obtained using a Keyence BZ-X710 microscope (Keyence, Osaka, Japan).

### Metabolite extraction and metabolome analysis^65^

After the culture medium was aspirated, plated HDF-75 cells were washed twice with PBS solution (10 mL and then 2 mL), treated with 740 µL of methanol, and left at rest for 30 s to inactivate enzymes. The cell extract was then treated with 500 µL of Milli-Q water containing internal standards (10 µM methionine sulfone) and left at rest for a further 30 s. Three hundred and fifty microliters of the extract was centrifugally filtered through a Millipore 5 kDa cutoff filter (UltrafreeMC-PLHCC, HMT) to remove macromolecules (9,100 × *g*, 4 °C, 120 min). The filtrate was centrifugally concentrated and re-suspended in 30 µL of acetonitrile/ Milli-Q water (1:1 v/v) for metabolome analysis at Infinity Lab (Yamagata, Japan). HPLC instrument modules were obtained from the 1290 Infinity line (Agilent Technologies): binary pumps (Model G7120A), an autosampler (G7167B), and a temperature-controlled column compartment (Model G7116B). A stock solution of 100 mM ammonium acetate was prepared by dissolving ammonium acetate (0.1 mol) in water, adjusting to pH 9.0 with ammonium hydroxide, and correcting the final volume to 1 L with water. Hydrophilic interaction chromatography (HILIC) was performed using a 100 mm × 2.1 mm i.d.

InfinityLab Poroshell 120 HILIC-Z column (Agilent Technologies). Solvent A was prepared by mixing 100 mL of the stock solution with 900 mL of water to yield a final ammonium acetate concentration of 10 mM (pH 9.0). Solvent B was prepared by mixing 100 mL of the stock solution with 900 mL of acetonitrile to yield a final concentration of 10 mM ammonium acetate (pH 9.0) in 80% acetonitrile. The deactivator additive (5 mM methylenediphosphonic acid) was spiked into the solvents for a final concentration of 5 μM for analysis. The flow rate was 0.3 mL/min, and the column temperature was set at 45 °C. A sample volume of 1 μL was injected into the column for each experiment. The gradient began initially at 100% B, decreased linearly to 90% at 0.5 min, decreased linearly again to 70% at 10 min, and subsequently re-equilibrated at 95% B for a further 3 min. The sample temperature was maintained at 4 °C in the autosampler prior to analysis. Mass spectrometry analysis was performed using an Agilent 6470 B triple quadrupole mass spectrometer (Agilent Technologies) equipped with a Jet Stream ESI probe in positive-ion mode. A capillary voltage of 3,500 V, a source temperature of 200 °C, gas flow of 10 L/min, and nebulizer of 40 psi, sheath gas temperature of 300 °C, sheath gas flow of 12 L/min were used. Raw data were sampled at 667 ms/cycle in Dynamic Multiple Reaction Monitoring (MRM) mode. The concentrations of individual nucleotides were calculated from the peak area in the chromatogram detected using MRM relative to the internal standard methionine sulfone.

### siRNA transfection

HDFs derived from the 75-year-old donor were transfected with AMPKα2 siRNA oligonucleotides (5’ ACCGAGCUAUGAAGCAGCUGGAUUU 3’) (Invitrogen, Waltham, MA, USA). Lipofectamine RNAiMAX (Thermo Fisher Scientific) was used for transfection according to the manufacturer’s instructions. For each well of 96-well plate, 1 pmol of siRNA oligonucleotides and 0.3 μl lipofectamin RNAiMAX was added into 100 μl culture media of the cells.

### Immunoblotting

Cells were lysed in RIPA buffer containing protease and phosphatase inhibitor cocktails (Nacalai Tesque, Kyoto, Japan). Cell lysates were sonicated for 10 s and heated at 95 °C for 5 min. Cell lysates were separated on 12 % or 4–20 % gels (Bio-Rad, Hercules, CA, United states) for SDS-PAGE. The proteins were transferred onto PDVF membranes (Bio-Rad) and blocked in blocking buffer (Nacalai Tesque) for 1 h. Next, the membranes were incubated overnight at 4 °C with the primary antibodies. The membranes were rinsed three times with TBS-T ((Tris-buffered saline)-0.01% Tween), then incubated with secondary HRP-conjugated antibodies for 1 h at 25 °C. The membrane was washed with TBS-T and treated with ImmunoStar Zeta reagent (Wako Pure Chemical Industries). Chemiluminescent images of the membranes were acquired using LuminoGraph I (WSE-6100; ATTO).

### Fatty acid oxidation (FAO) activity

FAO activity in HDFs derived from a 75-year-old donor was detected using FAOBlue (Funakoshi, Tokyo, Japan), according to the manufacturer’s instructions. HDFs were seeded at 5,000 cells/well in 96-well plates and treated with proAX (final concentration 100 μM) for 3, 6, and 24 h. The cells were then treated with FAO blue (final concentration 5 μM) for 2 h. The fluorescence (excitation/emission = 405/460 nm) of the cells was measured using an Infinite 200PRO M Plex microplate reader (Tecan, Zürich, Switzerland).

### Glucose consumption and lactate release

HDFs derived from 75-year-old donors were seeded at 5,000 cells/well in 96-well plates and treated with proAX (final concentration 100 μM) for 3, 6, and 24 h. The supernatant was collected and diluted 1:30 for the glucose assay or 1:10 for the lactate assay in ultrapure water. The diluted supernatant was measured using a glucose assay kit or glycolysis/OXPHOS assay kit (Dojindo, Kumamoto, Japan). Absorbance was measured at 450 nm using an Infinite 200PRO M Plex microplate reader (Tecan, Zürich, Switzerland).

### Extracellular flux analyzer

A Seahorse XF HS Mini Analyzer (Agilent Technologies) was used to measure OCR. HDFs derived from 75-year-old donors were seeded at 8,000 cells/well in XFp cell culture miniplates and treated with proAX (final concentration, 50 μM) for 3 and 24 h. Mitochondrial functions were examined through sequential injections of oligomycin (1.5 μM), Carbonyl cyanide-4 (trifluoromethoxy) phenylhydrazone (FCCP) (5 μM), and rotenone/antimycin A (Rot/AA) (0.5 μM). The obtained data were analyzed using the Wave software (version 10.0.1.72; Agilent Technologies).

### Oxidative stress assay in HDFs

The viability of HDFs was measured using the cell counting kit-8 (CCK-8) (Dojindo, Kumamoto, Japan). HDFs were seeded at 5,000 cells/well in 96-well plates. Treatment with proAX (final concentration 100 μM), HDFs were incubated for 24 h. After washing out the pretreatment medium, cells were exposed to H_2_O_2_ (300–400 μM) in serum-free medium (DMEM/F12) at 37 °C. After incubation for 2 h, the absorbance of the cells was measured at 450 nm using an Infinite 200PRO M Plex microplate reader (Tecan, Zürich, Switzerland).

### C. elegans cultures

Wild-type N2 and *aak-2* mutant *C. elegans* were provided by the Caenorhabditis Genetics Center (CGC, University of Minnesota). The *C. elegans* mutant strain RB754 (genotype: *aak-2 (ok524)* X) carries a loss-of-function mutation in *aak-2*, which encodes the *C. elegans* ortholog of AMPK^66^. The worms were grown and maintained on nematode growth medium (NGM) plates spread with live or dead *E. coli* OP50 as food. Dead OP50 was prepared by incubating in 65 °C for 30 min. All the worms were cultured at 25 °C. proAX was added to the NGM plate at final concentrations from 25 to 300 μM. For lifespan assays, a concentration of 100 μM was used for the *aak-2* mutant. elegans. The comparison samples were used in which 100 μM AMP, ATP, and AICAR were added to the NGM plate. All NGM plates were supplemented with DMSO at a final concentration of 1%.

### Lifespan Assays

After the age-synchronized worms were cultured to the L4 stage, 35 worms were transferred to the desired NGM plates. The worms were transferred to fresh NGM plates with or without drugs every 1–2 days. Plates without any added drugs were used as the control. Live and dead worms were counted during transfer. Worms that displayed no movement after gentle contact with a platinum picker were considered dead. Kaplan–Meier curves were generated using Graph Pad Prism 10 (GraphPad Software, CA, USA) and calculated by Excel (Windows 365).

### Body length of wild-type N2 C. elegans

After age synchronization, the worms were transferred to the desired NGM plates and cultured for 72 h. The body length of each live worm was measured using a VHX-900F instrument (KEYENCE, Osaka, Japan).

### ATP levels of wild-type N2 C. elegans

After age synchronization, the worms were transferred to the NGM plates with or without proAX and cultured for 4 days. Bacteria attached to the bodies of 7-day-old worms were removed by allowing the worms to crawl onto an NGM agar plate with M9 buffer. Worms were collected in M9 buffer, and endogenous ATP was extracted by sonication on ice for 1 min (Ultrasonic Homogenizer UP-21P; TOMY, Tokyo, Japan) and centrifugation (15,000 rpm, 10 min, 4 °C). An ATP assay kit (Dojindo, Kumamoto, Japan) was used to measure the endogenous ATP levels. Luminescence was measured using an Infinite 200PRO M Plex microplate reader (Tecan).

### Paraquat sensitivity of wild-type N2 C. elegans

After age synchronization, the worms were transferred to the NGM plates with or without proAX and cultured for 4 days. To assay paraquat sensitivity, the 7-day-old worms were collected and transferred to NGM plates containing 200 mM paraquat. Worms that displayed no movement after gentle contact with a platinum picker were considered dead.

### Measurement of reactive oxygen species of wild-type N2 C. elegans

After age synchronization, worms were transferred to NGM plates with or without proAX and cultured for 4 days. On day 7, bacteria adhered to the worm surface were removed by allowing the animals to crawl onto NGM agar plates containing M9 buffer. Worms were then collected in M9 buffer, and endogenous ROS was extracted by sonication on ice for 30 s using an ultrasonic homogenizer (UP-21P; TOMY, Tokyo, Japan), followed by centrifugation at 15,000 rpm for 10 min at 4 °C. ROS levels were measured using a DCFH-DA (Med Chem Express, NJ, USA). Fluorescence was quantified with an Infinite 200PRO M Plex microplate reader (Tecan).

### Statistics and reproducibility

Data are presented as mean ± s.d. for the indicated number of biological replicates (n) from least three independent experiments. Statistical analyses were performed using GraphPad Prism v.10.2 (GraphPad Software, CA, USA). The relevant statistical tests are described in the figure legends. Time-to-event data were analyzed using Kaplan–Meier curves, and differences in survival were determined using the log-rank test by Excel (for Microsoft 365). P set at *p* < 0.05 was significant.

## Supporting information

Supporting Infomation

## Acknowledgements

**General:** We thank to Takamasa Ishikawa and Ryuhei Kudo (Infinity Lab, INC, Yamagata, Japan) for metabolome analysis and measurement of AXP.

**Funding:** This study was supported in part by Grants-in-Aid for Scientific Research (KAKENHI) Grant Number JP23K18580 (for T. A.), JP19H05720 (for M. T.), JP22K19759 (for E. N. K.) from the Japan Society for the Promotion of Science (JSPS). Additionally, this study was partially supported by the Dynamic Alliance Open Innovation Bridging Human, Environment and Materials from the Ministry of Education, Culture, Sports, and Technology of Japan (MEXT) (for M. T., and T. A.), CORE^2^-A lab (for T. A.). This study was partially supported by AMED-Hashiwatashi research program A (for T. A.), JST SPRING Japan Grand Number JPMJSP2136 (for M. K.) and Nagase Science and Technology Foundation (for E. N. K.).

## Author Information

### Corresponding Authors

T. Anada, M. Tanaka

Institute for Materials Chemistry and Engineering, Kyushu university, 744 Motooka, Nishi-ku Fukuoka 819-0395, Japan

### Authors

T. Anada, S. Kobayashi, M. Tanaka

Institute for Materials Chemistry and Engineering, Kyushu university, 744 Motooka, Nishi-ku Fukuoka 819-0395, Japan

T. Anada, M. Kawahara, T. Shimada, R. Kuroda, M. Tanaka

Department of Applied Chemistry, Graduate School of Engineering, Kyushu University, 744 Motooka, Nishi-ku Fukuoka 819-0395, Japan

H. Okamura, F. Nagatsugi

Institute of Multidisciplinary Research for Advanced Materials, Tohoku University, 2-1-1 Katahira, Aoba-ku, Sendai, 980-8577 Japan

1. H. Okamura

Faculty of Medicine, Dentistry and Pharmaceutical Sciences, Okayama University, 1-1-1 Tsushima-naka, Kita-ku, Okayama 700-8530, Japan

D. Setoyama, Y. Kunisaki

Department of Clinical Chemistry and Laboratory Medicine, Kyushu University Hospital, Fukuoka 812-8582, Japan

E. Kage-Nakadai

Laboratory of Environment and Health Science, Graduate School of Human Life and Ecology, Osaka Metropolitan University, 3-3-138 Sugimoto, Sumiyoshi-ku, Osaka 558-8585, Japan

E. Kage-Nakadai

Institute for Life and Medical Sciences, Kyoto University, 53 Shogoin Kawahara-cho, Sakyo-ku, Kyoto 606-8507, Japan

