## Supporting Infomation for "A nucleic acid prodrug that activates mitochondrial respiration, promotes stress resilience, and prolongs lifespan"


**This PDF includes:**
Figures S1 to S5
Tables S1 to S2
General Information for Synthesis

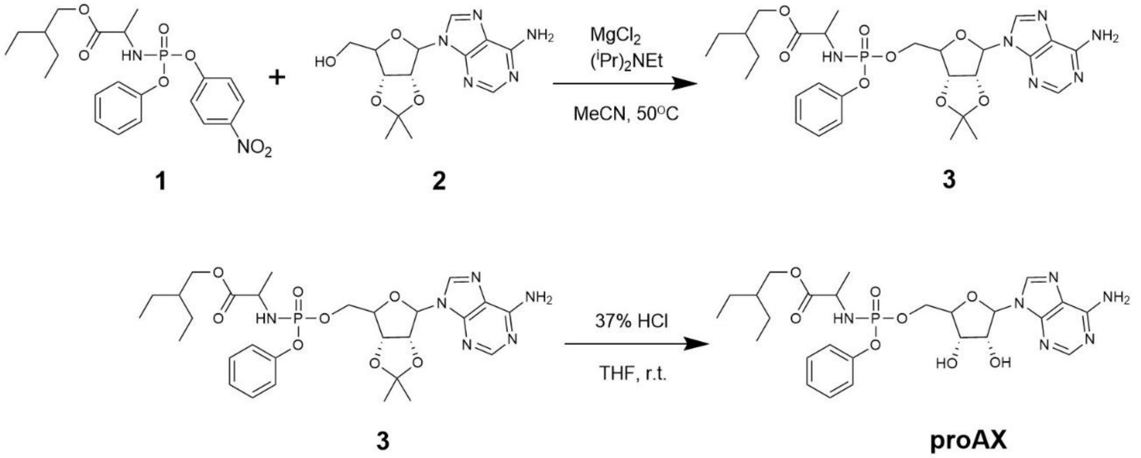


**Figure. S1**: Synthetic scheme for proAX. The experimental procedures and product characterization are described in the Supporting Information.

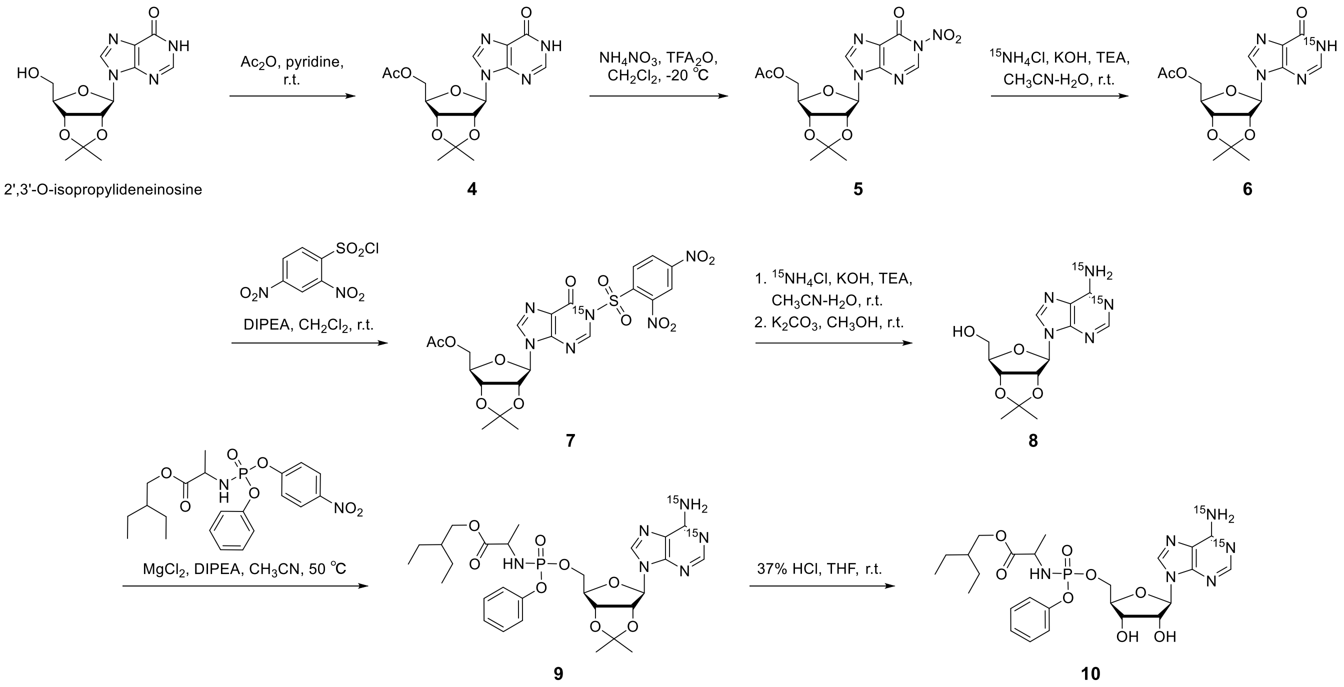


**Figure. S2**: Synthetic scheme for [$^{15}N_2$]-proAX (**10**). The experimental procedures and product characterization are described in Supporting Information.

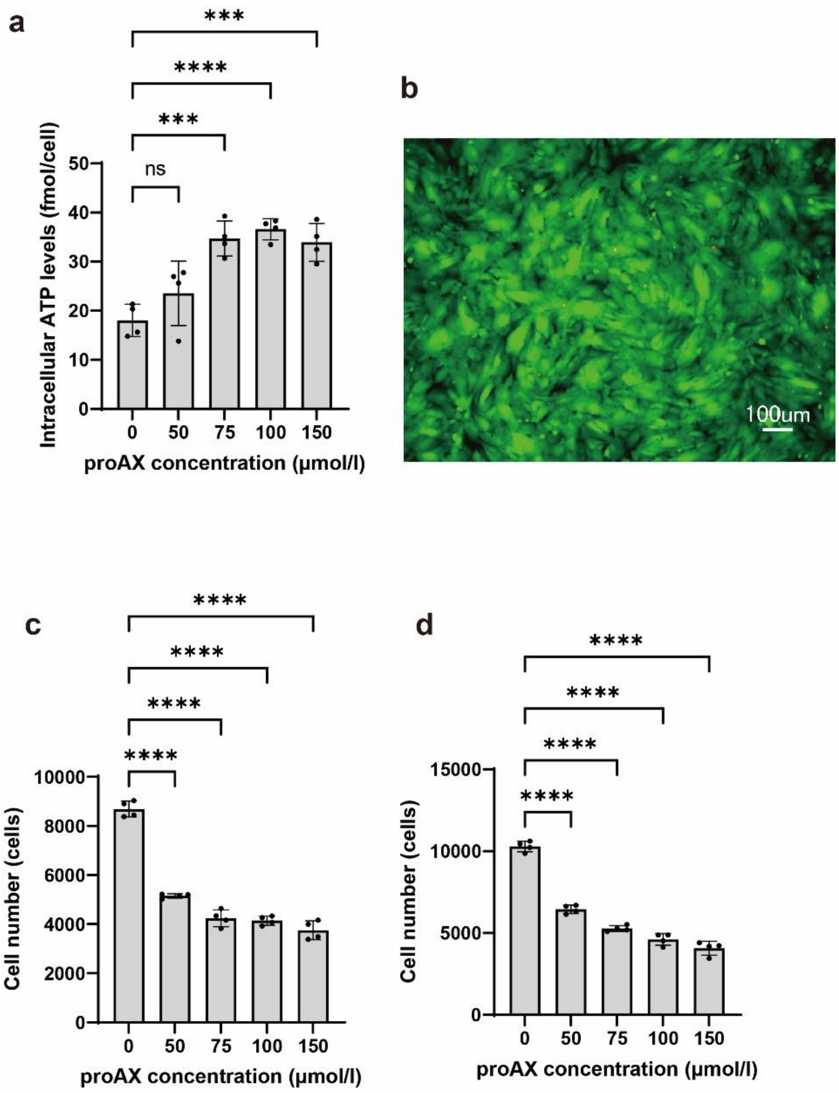


**Figure. S3**: **a,** ProAX enhances intracellular ATP levels in HDF-29 cells. **b**, Live (green) and dead (red) viability analysis for the evaluation of proAX cytotoxicity in HDF-75 cells. Effect of proAX on cell proliferation in HDF-75 (**c**), and HDF-29 (**d**). For panels a, c, and d, mean ± standard deviation shown, along with statistical significances for relevant comparisons. Significance is assessed using a one-way ANOVA with Dunnett's multiple comparison test, with *n* five replicates per condition. ****$p < 0.0001$, ***$p < 0.001$; ns indicates not significant.

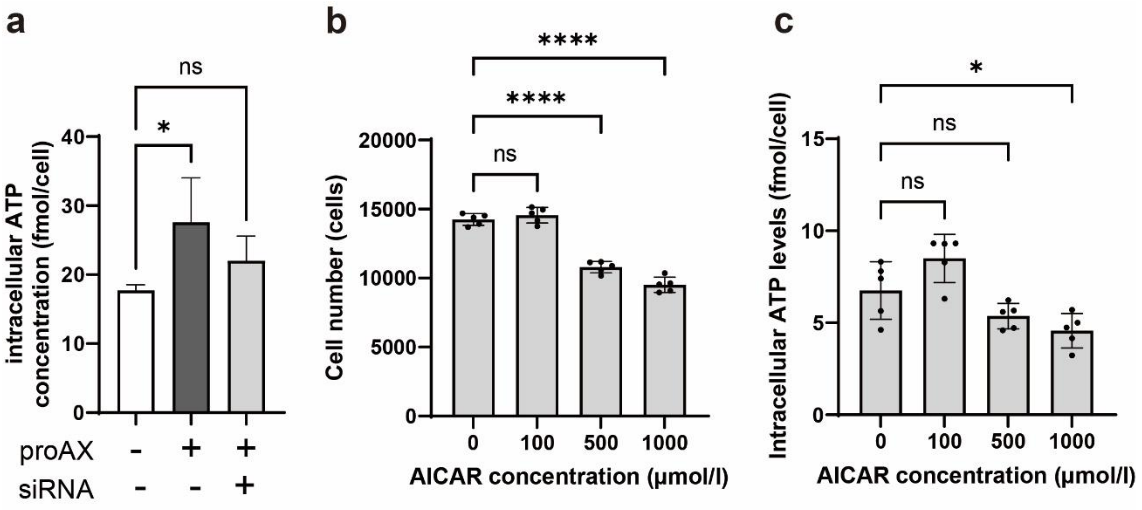


**Figure. S4**: **a**, Intracellular ATP changes in HDF-75 treated with 100 μM proAX in the presence or absence of AMPKa2 siRNA. **b,** AICAR did not increase intracellular ATP levels in HDF-75 cells. **c,** AICAR suppresses the proliferation of HDF-75 cells. Mean ± standard deviation shown, along with statistical significances for relevant comparisons. Significance is assessed using a one-way ANOVA with Dunnett's multiple comparison test, with $n$ five replicates per condition. ****$p < 0.0001$, *$p < 0.05$; ns indicates not significant.

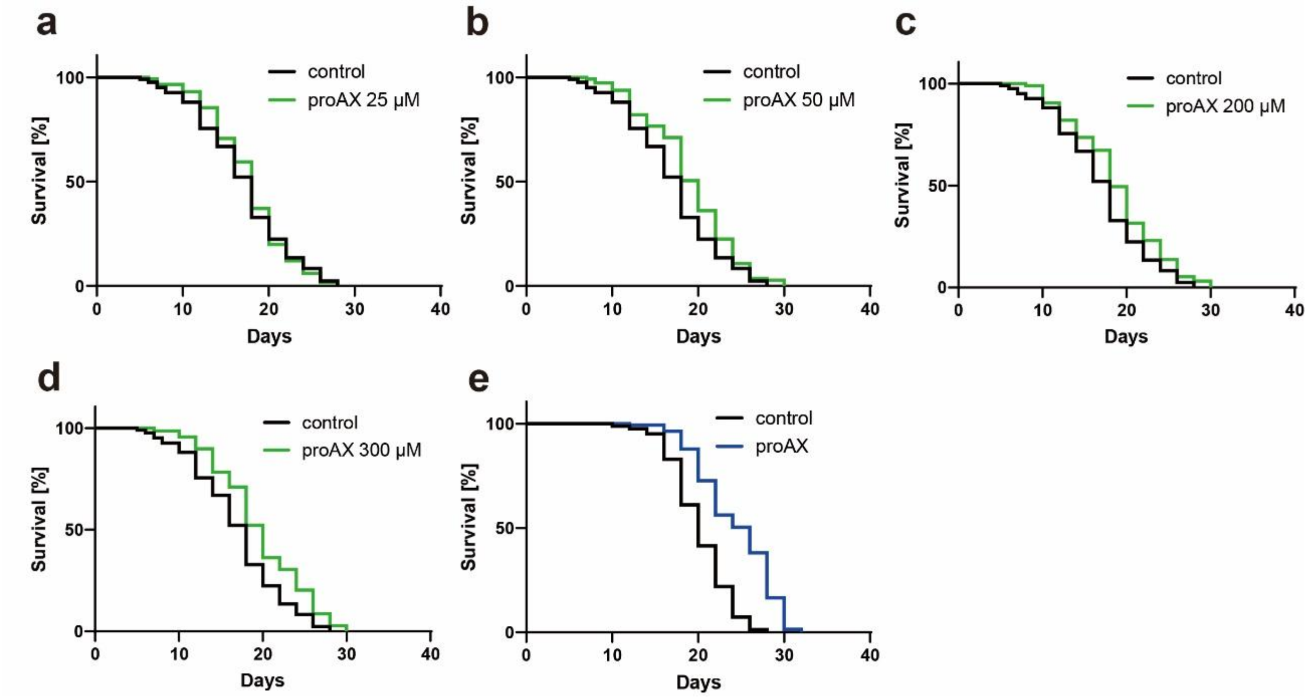


**Figure. S5**: ProAX extends the lifespan of *Caenorhabditis*. *elegans*. **a-d**, Survival curves of *C. elegans* treated with proAX (0–300 μM). **e**, Survival curves of *C. elegans* treated with proAX (100 μM) with feeding dead *Escherichia coli*. Significance is calculated using the log-rank test.

**Table S1**. Effect of proAX on lifespan extension of *C. elegans* (feeding on dead *E. coli*).

| **sample** | **Mean lifespan ± SE (days)** | **Extension rate (%)** | **N** | ***p* value (log-rank)** |
|---|---|---|---|---|
| Control (no drug) | 20.6 ± 0.4 | - | 82 | - |
| 100 μM proAX | 24.9 ± 0.4 | 20.9 | 139 | $< 10^{-4}$ |

**Table S2**. Effect of proAX on lifespan extension of *aak-2* mutant worms.

| sample | Mean lifespan ± SE (days) | Extension rate (%) | N | *p* value (log-rank) |
|---|---|---|---|---|
| Control (no drug) | 16.4 ± 0.3 | - | 87 | - |
| 100 μM proAX | 17.3 ± 0.4 | 5.5 | 87 | 0.1014 |

### General Information for Synthesis of proAX

(*S*)-2-Ethylbuthyl 2-{[(*S*)-(4-nitrophenoxy)(phenoxy)phosphoryl]amino}propanoate (**1**) was purchased from combi-blocks (San Diego, CA, United States). 2',3'-*O*-Isopropylideneadenosine (**2**) was obtained from TCI (Tokyo, Japan). Other chemical reagents and solvents, such as magnesium chloride, *N,N*-diisopropylethylamine, were commercially available. Nuclear magnetic resonance (NMR) spectra were recorded on a Bruker AVANCE III 400 MHz at room temperature (25 ºC), with tetramethylsilane as an internal standard. Proton nuclear magnetic resonance spectra are reported in parts per million (ppm) on the δ scale and are referenced from the residual protium in the NMR solvent (methanol-$d_4$: δ 3.31, water-$d_2$: δ 4.79). Data is reported as follows: chemical shift [multiplicity (s = singlet, d = doublet, t = triplet, q = quartet, m = multiplet, br = broad), coupling constants (*J*) in Hertz, integration. Carbon-13 nuclear magnetic resonance spectra are reported in parts per million on the δ scale and are referenced from the carbon resonances of the solvent (methanol-$d_4$: δ 49.00). Data is reported as follows: chemical shift. Phosphorus-31 nuclear magnetic resonance spectra are reported in parts per million on the δ scale. Data is reported as follows: chemical shift. Fast atom bombardment mass spectra (FAB-MS) were measured on double-focusing mass spectrometer (JEOL, JMS-700T).

### Experimental Procedures and Product Characterization

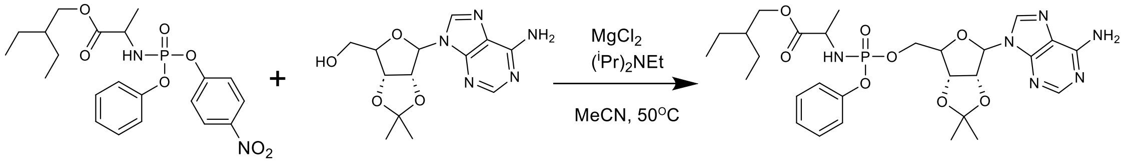


1 2 3

#### (*S*)-2-Ethylbutyl 2-{[(*S*)-{[(2*R*,3*S*,4*R*,5*R*)-5-(6-amino-9*H*-purin-9-yl)-2,2-dimethyltetrahydrofuro(3.4-*d*)(1,3)dioxol-4-yl]methoxy}(phenoxy)phosphoryl]amino}propanoate (3)

(*S*)-2-Ethylbuthyl 2-{[(*S*)-(4-nitrophenoxy)(phenoxy)phosphoryl]amino}propanoate (**1**, 3.58 g, 7.95 mmol, 1.20 equiv), 2',3'-*O*-isopropylideneadenosine (**2**, 2.04 g, 6.64 mmol, 1 equiv)), magnesium chloride (632 mg, 6.64 mmol, 1.00 equiv) were dissolved in acetonitrile (32 mL) at room temperature, and the solution was heated at 50 ºC for 10 min. *N,N*-Diisopropylethylamine (2.90 mL, 16.60 mmol, 2.50 equiv) was added and allowed to react for 20 min. After cooling the reaction mixture to room temperature, diluted with ethyl acetate (200 mL). The organic layer was washed with 5% aqueous citric acid solution (80 mL), saturated aqueous ammonium chloride solution (80 mL), 5% aqueous potassium carbonate solution (2 × 80 mL), and brine (80 mL), and dried over anhydrous sodium sulfate, and evaporated. The obtained concentrate was subjected to silica gel chromatography eluting with 0 - 50% hexane in acetones to afford the product **3** (3.87 g, 94%). $^{1}$H NMR (400 MHz, methanol-$d_4$) : δ 8.23 (s, 1H), 8.20 (s, 1H), 7.36 – 7.09 (m, 5H), 6.21 (d, *J* = 2.4 Hz, 1H), 5.38 (dd, *J* = 2.4, 6.2 Hz 1H), 5.11 (dd, *J* = 6.3, 3.2 Hz, 1H), 4.44 (br, 1H), 4.36 – 4.19 (m, 2H), 4.06 – 3.91 (dd, *J* = 6.0, 10.8 Hz, 2H), 3.88 (dd, *J* = 7.6, 9.6, 1H), 1.60 (s, 3H), 1.53 – 1.40 (m, 1H), 1.40 – 1.21 (m, 10H), 0.86 (t, *J* = 7.5 Hz, 6H).

**(*S*)-2-Ethylbutyl 2-{[(*S*)-{[(2*R*,3*S*,4*R*,5*R*)-5-(6-amino-9*H*-purin-9-yl)-3,4-dihydroxytetrahydrofuran-2-yl]methoxy}(phenoxy)phosphoryl]amino}propanoate (proAX)**

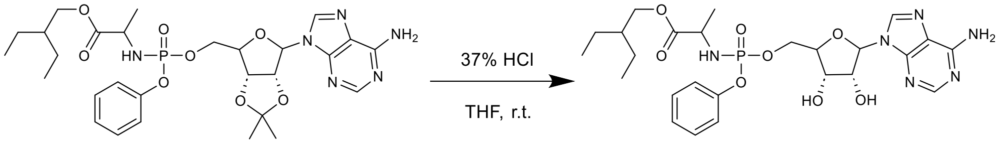


**3** **proAX**

(*S*)-2-Ethylbutyl 2-{[(*S*)-{[(2*R*,3*S*,4*R*,5*R*)-5-(6-amino-9*H*-purin-9-yl)-2,2-dimethyltetrahydrofuro(3.4-*d*)(1,3)dioxol-4-yl]methoxy}(phenoxy)phosphoryl]amino}propanoate (**3**, 3 g) was dissolved in tetrahydrofuran (30 mL). The solution was added 37% aqueous hydrochloric acid solution (6 mL) slowly at 0 ºC, and the reaction mixture was allowed to warm to room temperature, and stirred for 5 h. After that, diluted with water (30 mL) and adjusted to pH = 8 by the addition of saturated aqueous sodium bicarbonate solution (60 mL), and extracted with ethyl acetate (30 mL). The organic layer was washed with brine (15 mL), and dried over anhydrous sodium sulfate, and evaporated. The obtained concentrate was subjected to silica gel chromatography eluting with solution containing 80% ethyl acetate and 20% methanol to afford proAX (0.78 g, 26%). $^{1}$H NMR (400 MHz, methanol-*d*$_4$) : δ 8.24 (s, 1H), 8.20 (s, 1H), 7.38 – 7.13 (m, 5H), 6.03 (d, *J* = 4.9 Hz, 1H), 4.63 (t, *J* = 5.1 Hz, 1H), 4.39 (m, 2H), 4.32 (m, 1H), 4.25 (br, 1H), 4.02 (dd, *J* = 10.9, 5.7 Hz, 1H), 3.93 (m, 2H), 1.44 (q, *J* = 6.1 Hz, 1H), 1.32 (dd, *J* = 8.2, 6.1 Hz, 7H), 0.85 (t, *J* = 7.4 Hz, 6H) ; $^{13}$C NMR (101 MHz, methanol-*d*$_4$) δ 174.98, 174.93, 156.55, 152.70, 152.17, 152.10, 150.61, 141.47, 130.76, 126.16, 126.15, 121.38, 121.33, 120.55, 90.12, 84.47, 84.39, 75.44, 71.61, 68.20, 67.60, 67.55, 51.61, 51.60, 49.71, 49.64, 49.50, 49.43, 49.28, 49.21, 49.07, 49.00, 48.86, 48.79, 48.57, 48.36, 41.75, 24.25, 24.23, 20.62, 20.55, 16.26, 11.31, 11.27, -0.06 ; $^{31}$P NMR (162 MHz, methanol-*d*$_4$) δ 4.86 ; FAB-MS (m/z): [M]+ calcd for $C_{25}H_{35}N_6O_8P$, 578.6; found, 579.2.

**General Information for Synthesis of [$^{15}N_2$]-proAX**

(*S*)-2-Ethylbuthyl 2-{[(*S*)-(4-nitrophenoxy)(phenoxy)phosphoryl]amino}propanoate (**1**) was purchased from combi-blocks (San Diego, CA, United States). $^{15}NH_4Cl$ (≥98 atom% $^{15}$N) was purchased from Sigma-Aldrich. All other reagents were purchased from Tokyo Chemical Industry, Merck, FUJIFILM Wako Pure Chemical Corporation, Kanto Chemical and Thermo Fisher Scientific. Nuclear magnetic resonance (NMR) spectra were recorded on a Bruker AVANCE III 400 MHz and 600 MHz at 298 K. $^{1}$H NMR spectra are reported in parts per million (ppm) on the δ scale and are referenced from the residual protium in the NMR solvent (methanol-*d*4: δ 3.31, water-*d*2: δ 4.79, DMSO-*d*6: δ 2.50). Data is reported as follows: chemical shift [multiplicity (s = singlet, d = doublet, t = triplet, q = quartet, m = multiplet, br = broad), coupling constants (*J*) in Hertz, integration. $^{13}$H NMR spectra are reported in parts per million on the δ scale and are referenced from the carbon resonances of the solvent (methanol-*d*4: δ 49.0, DMSO-*d*6: δ 39.5). Data is reported as follows: chemical shift. Phosphorus-31 nuclear magnetic resonance spectra are reported in parts per million on the δ scale. Data is reported as follows: chemical shift. Fast atom bombardment mass spectra (FAB-MS) were measured on double-focusing mass spectrometer (JEOL, JMS-700T), and electron spray ionization mass spectra (ESI-MS) were measured on MicrOTOF-QII (Bruker).

**5'-Acetyl-2',3'-*O*-(1-methylethylidene)inosine (4)**

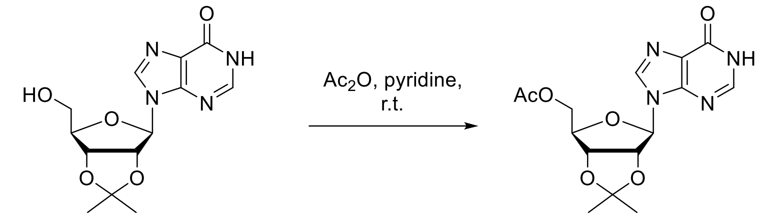


2',3'-O-isopropylideneinosine **4**

To a solution of 2',3'-*O*-isopropylideneinosine (5.3 g, 17.2 mmol) in dry pyridine (86 mL) was added acetic anhydride (3.3 mL, 34.4 mmol) and DMAP (10 mg), and the reaction mixture was stirred at room temperature for 1 h. The solvent was evaporated, and the residue was diluted with $CH_2Cl_2$. The organic layer was washed with water, 0.5 M aqueous hydrochloric acid solution, saturated aqueous $NaHCO_3$ solution and brine, and dried over $Na_2SO_4$. The solvent was evaporated to afford compound **4** (4.46 g, 12.7 mmol, 74%) as a white solid. $^1$H NMR (600 MHz, DMSO-$d_6$) δ 12.46 (br, 1H), 8.24 (s, 1H), 8.09 (s, 1H), 6.16 (d, *J*= 2.4 Hz, 1H), 5.38 (dd, *J*= 6.2, 2.4 Hz, 1H), 5.00 (dd, *J*= 6.2, 3.3 Hz, 1H), 4.44-4.42 (m, 1H), 4.20 (dd, *J*= 11.8, 4.4 Hz, 1H), 4.13 (dd, *J*= 11.8, 6.2 Hz, 1H), 1.96 (s, 3H), 1.54 (s, 3H), 1.33 (s, 3H); $^{13}$C NMR (151 MHz, DMSO-$d_6$) δ 170.03, 156.53, 147.70, 146.13, 138.95, 124.66, 113.52, 89.22, 83.75, 83.57, 80.97, 63.70, 26.96, 25.21, 20.48; ESI-MS (m/z): $[M+Na]^+$ calcd for $C_{15}H_{18}N_4NaO_6^+$, 373.1119; found, 373.1115.

**5'-Acetyl-2',3'-*O*-(1-methylethylidene)-1-nitroinosine (5)**

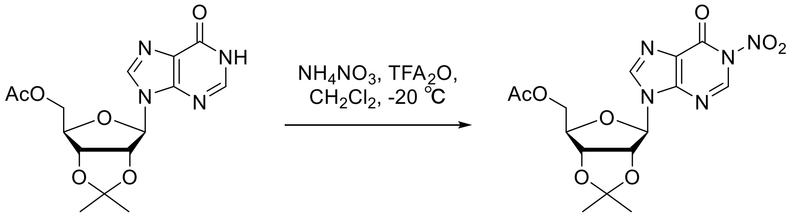


**4** **5**

To a suspension of ammonium nitrate (5 g, 62.5 mmol) in dry $CH_2Cl_2$ (100 mL) was added trifluoroacetic anhydride (18 mL, 129 mmol), and the suspension was stirred at room temperature for 1 h. The resulting yellow solution was cooled to -25 ºC, and compound **4** (2.7 g, 7.71 mmol) was added portionwise. The reaction mixture was stirred at the same temperature for 1.5 h. The reaction mixture was diluted with 0.5 M aqueous triethylammonium acetate buffer (pH 7.0) and extracted with $CH_2Cl_2$. The organic layer was washed with brine and dried over $Na_2SO_4$. The solvent was evaporated, and the residue was purified by silica gel column chromatography ($CH_2Cl_2$:MeOH = 100:0 to 96:4) to afford compound **5** (2.15 g, 5.44 mmol, 71%) as a yellow foam. $^1$H NMR (600 MHz, DMSO-$d_6$) δ 9.11 (s, 1H), 8.24 (s, 1H), 8.43 (s, 1H), 6.19 (d, *J*= 2.2 Hz, 1H), 5.36 (dd, *J*= 6.2, 2.2 Hz, 1H), 5.00 (dd, *J*= 6.2, 3.2 Hz, 1H), 4.44-4.42 (m, 1H), 4.20 (dd, *J*= 12.0, 4.4 Hz, 1H), 4.16 (dd, *J*= 12.0, 6.0 Hz, 1H), 1.95 (s, 3H), 1.54 (s, 3H), 1.33 (s, 3H); $^{13}$C NMR (151 MHz, DMSO-$d_6$) δ 170.01, 149.65, 145.15, 143.04, 140.80, 123.87, 113.57, 88.69, 84.16, 83.85, 80.88, 63.67, 26.90, 25.17, 20.48; ESI-MS (m/z): $[M+Na]^+$ calcd for $C_{15}H_{17}N_5NaO_8^+$, 418.0969; found, 418.0971.

**5'-Acetyl-2',3'-*O*-(1-methylethylidene)-[1-$^{15}$N]inosine (6)**

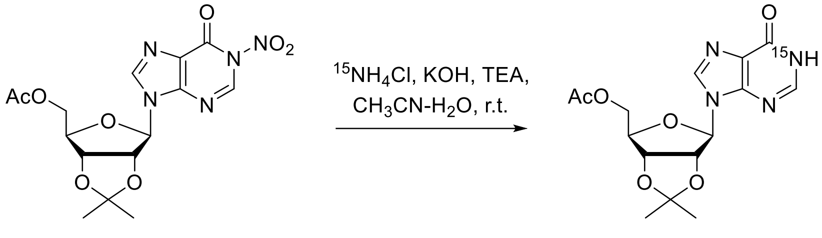


5 6

To a solution of $^{15}NH_4Cl$ (347 mg, 6.37 mmol), 1 M potassium hydroxide (6.64 mL, 6.64 mmol) and triethylamine (773 µL, 5.58 mmol) in water (23 mL) was added a solution of compound **5** (2.1 g, 5.31 mmol) in $CH_3CN$ (86 mL), and the mixture was stirred at room temperature for 2 h. The solvent was evaporated, and the residue was diluted with $CH_2Cl_2$. The organic layer was washed with 5% aqueous citric acid solution and brine, and dried over $Na_2SO_4$. The solvent was evaporated, and the residue was purified by silica gel column chromatography ($CH_2Cl_2$:MeOH = 100:0 to 94:6) to afford compound **6** (1.44 g, 4.10 mmol, 77%) as a pale yellow powder. $^1$H NMR (600 MHz, DMSO-*d*$_6$) δ 12.4 (dd, $J_{HN}$= 89.6, 4.0 Hz, 1H), 8.25 (s, 1H), 8.09 (dd, $J_{HN}$= 7.4, 4.0 Hz, 1H), 6.16 (d, *J*= 2.4 Hz, 1H), 5.38 (dd, *J*= 6.2, 2.4 Hz, 1H), 5.00 (dd, *J*= 6.2, 3.2 Hz, 1H), 4.44-4.42 (m, 1H), 4.20 (dd, *J*= 11.8, 4.4 Hz, 1H), 4.13 (dd, *J*= 11.8, 6.2 Hz, 1H), 1.96 (s, 3H), 1.54 (s, 3H), 1.33 (s, 3H); $^{13}$C NMR (151 MHz, DMSO-*d*$_6$) δ 170.04 156.50 (d, $J_{CN}$= 10.6 Hz), 147.70 (d, $J_{CN}$= 3.0 Hz), 146.10 (d, $J_{CN}$= 9.1 Hz), 138.96, 124.65 (d, $J_{CN}$= 7.6 Hz), 113.52, 89.22, 83.75, 83.57, 80.97, 63.70, 26.96, 25.21, 20.48; ESI-MS (m/z): [M+Na]$^+$ calcd for $C_{15}H_{18}N_3{}^{15}NNaO_6^+$, 374.1089; found, 374.1094.

**5'-Acetyl-2',3'-*O*-(1-methylethylidene)-1-[(2,4-dinitrophenyl)sulfonyl]-[1-$^{15}$N]inosine (7)**

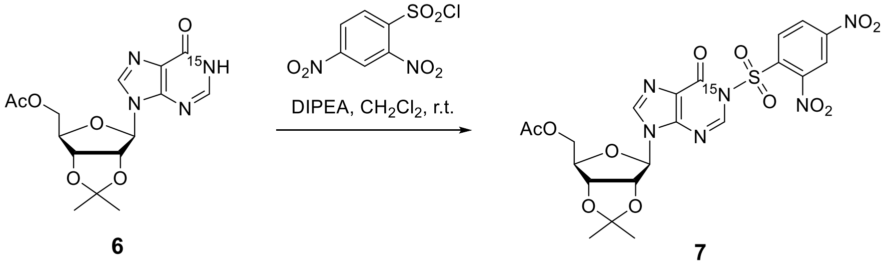


6 7

To a solution of compound **6** (1.4 g, 3.98 mmol) in $CH_2Cl_2$ was added *N*,*N*-diisopropylethylamine (905 µL, 5.18 mmol) and 2,4-dinitrobenzenesulfonyl chloride (1.53 g, 4.78 mmol), and the mixture was stirred at room temperature for 1.5 h. The reaction mixture was diluted with EtOAc. The organic layer was washed with water and brine, and dried over $Na_2SO_4$. The solvent was evaporated, and the residue was purified by silica gel column chromatography (Hexane:EtOAc = 80:20 to 40:60) to afford compound **7** (1.72 g, 2.96 mmol, 74%) as a yellow powder. $^1$H NMR (600 MHz, DMSO-*d*$_6$) δ 8.97-8.96 (m, 1H), 8.79-8.78 (m, 2H), 8.77 (d, $J_{HN}$= 6.6 Hz, 1H), 8.41 (s, 1H), 6.22 (d, *J*= 2.2 Hz, 1H), 5.38 (dd, *J*= 6.2, 2.2 Hz, 1H), 5.03 (dd, *J*= 6.2, 3.2 Hz, 1H), 4.45-4.43 (m, 1H), 4.20 (dd, *J*= 12.0, 4.4 Hz, 1H), 4.16 (dd, *J*= 12.0, 6.0 Hz, 1H), 1.94 (s, 3H), 1.55 (s, 3H), 1.34 (s, 3H); $^{13}$C NMR (151 MHz, DMSO-*d*$_6$) δ 169.98, 153.29 (d, $J_{CN}$= 4.5 Hz), 151.55, 147.83, 146.72, 143.97 (d, $J_{CN}$= 3.0 Hz), 140.89, 136.67, 132.83 (d, $J_{CN}$= 4.5 Hz), 127.50, 123.35 (d, $J_{CN}$= 9.1 Hz), 121.18, 113.59, 89.72, 84.19, 83.85, 80.88, 63.64, 26.92, 25.22, 20.45; ESI-MS (m/z): [M+Na]$^+$ calcd for $C_{21}H_{21}N_5{}^{15}NO_{12}S^+$, 582.0903; found, 582.0900.

### 2',3'-*O*-Isopropylidene-[1,6-$^{15}$N]adenosine (8)

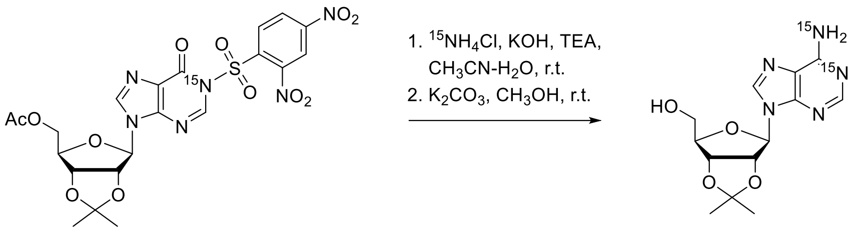


**7** **8**

To a solution of $^{15}NH_4Cl$ (191 mg, 3.51 mmol), 1 M potassium hydroxide (3.51 mL, 3.51 mmol) and triethylamine (480 µL, 3.51 mmol) in water (12 mL) was added a solution of compound **7** (1.7 g, 2,92 mmol) in $CH_3CN$ (48 mL), and the mixture was stirred at r.t. for 2.5 h followed by refluxing for 30 min. The solvent was evaporated, and the residue was diluted with EtOAc. The organic layer was washed with saturated aqueous $NaHCO_3$ solution and brine, and dried over $Na_2SO_4$. The solvent was evaporated, and the residue was subjected to silica gel column chromatography ($CH_2Cl_2$:MeOH = 100:0 to 96:4). The appropriate fractions were combined and evaporated. The residue was dissolved in dry MeOH, and $K_2CO_3$ was added. The reaction mixture was stirred at room temperature for 30 min. The solvent was evaporated, and the residue was purified by silica gel column chromatography ($CH_2Cl_2$:MeOH = 100:0 to 96:4) to afford compound **8** (413 mg, 1.33 mmol, 46% over 2 steps) as an off-white powder. $^1$H NMR (600 MHz, DMSO-$d_6$) δ 8.34 (s, 1H), 8.15 (d, $J_{HN}$= 15.8 Hz, 1H), 7.35 (d, $J_{HN}$= 90.0 Hz, 2H), 6.12 (d, *J*= 3.0 Hz, 1H), 5.34 (dd, *J*= 6.0, 3.0 Hz, 1H), 5.24 (t, *J*= 5.6, 3.0 Hz, 1H), 4.96 (dd, *J*= 6.0, 2.4 Hz, 1H), 4.22-4.20 (m, 1H), 3.58-3.50 (m, 2H), 1.54 (s, 3H), 1.32 (s, 3H); $^{13}$C NMR (151 MHz, DMSO-$d_6$) δ 156.59 (dd, $J_{CN}$= 19.6, 4.5 Hz), 153.10, 149.28, 140.16, 119.56, 113.5190.08, 86.81, 83.68, 81.83, 62.05, 27.55, 25.66; ESI-MS (m/z): [M+Na]$^+$ calcd for $C_{13}H_{18}N_3{}^{15}N_2O_4{}^+$, 310.1294; found, 310.1291.

### 2-Ethylbutyl ((((3*aR*,4*R*,6*R*,6*aR*)-6-(6-amino-9*H*-[1,6-$^{15}$N]-purin-9-yl)-2,2-dimethyltetrahydrofuro[3,4-d][1,3]dioxol-4-yl)methoxy)(phenoxy)phosphoryl)-L-alaninate (9)

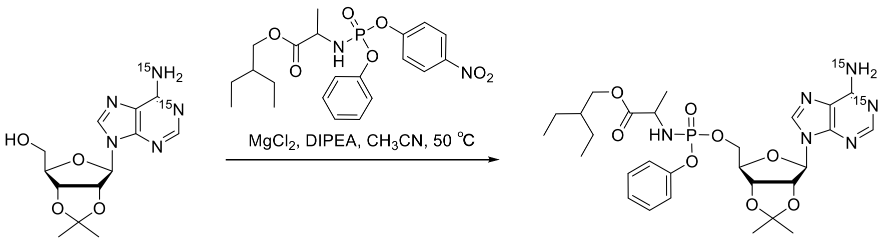


**8** **9**

(*S*)-2-Ethylbuthyl 2-{[(*S*)-(4-nitrophenoxy)(phenoxy)phosphoryl]amino}propanoate (**1**, 0.31 g, 0.37 mmol, 1.20 equiv), 2',3'-*O*-Isopropylidene-[1,6-$^{15}$N]adenosine (**8**, 0.18 g, 0.58 mmol, 1 equiv)), magnesium chloride (55 mg, 0.58 mmol, 1.00 equiv) were dissolved in acetonitrile (3 mL) at room temperature, and the solution was heated at 50 ºC for 10 min. *N*,*N*-Diisopropylethylamine (0.25 mL, 1.45 mmol, 2.50 equiv) was added and allowed to react for 20 min. After cooling the reaction mixture to room temperature, diluted with ethyl acetate (20 mL). The organic layer was washed with 5% aqueous citric acid solution (10 mL), saturated aqueous ammonium chloride solution (10 mL), 5% aqueous potassium carbonate solution (2 × 10 mL), and brine (10 mL), and dried over anhydrous sodium sulfate, and evaporated. The obtained concentrate was subjected to silica gel chromatography eluting with 0 - 50% hexane in acetones to afford the product **9** (0.14 g, 78%). $^1$H NMR (600 MHz, methanol-$d_4$) δ 8.24 (s, 1H), 8.20 (d, *J* = 15.3 Hz, 1H), 7.34 – 7.13 (m, 5H), 6.21 (d, *J* = 2.6 Hz, 1H), 5.38 (dd, *J* = 6.3, 2.5 Hz, 1H), 5.11 (dd, *J* = 6.3, 3.2 Hz, 1H), 4.44 (br, 1H), 4.35 – 4.22 (m, 2H), 4.04 – 3.93 (m, 2H), 3.92 – 3.87 (m, 1H), 1.59 (s, 3H), 1.48 – 1.43 (m, 1H), 1.36 – 1.24 (m, 10H), 0.86 (t, *J* = 7.5 Hz, 6H).

**2-Ethylbutyl ((((2*R*,3*S*,4*R*,5a*R*)-5-(6-amino-9*H*-[1,6-$^{15}$N]-purin-9-yl)-3,4-dihydroxytetrahydrofuran-2-yl)methoxy)(phenoxy)phosphoryl)-L-alaninate (10)**

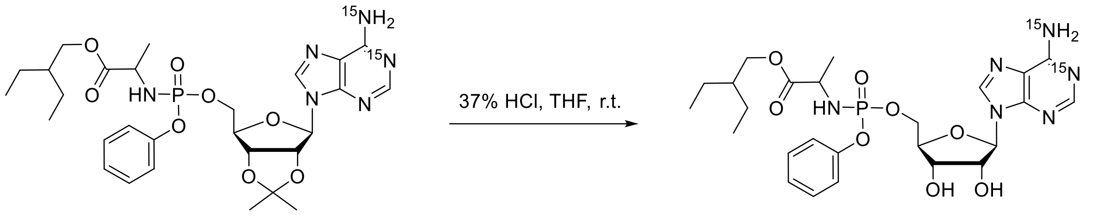


**9** **10**

2-Ethylbutyl ((((3*aR*,4*R*,6*R*,6*aR*)-6-(6-amino-9*H*-[1,6-$^{15}$N]-purin-9-yl)-2,2-dimethyltetrahydrofuro[3,4-d][1,3]dioxol-4-yl)methoxy)(phenoxy)phosphoryl)-L-alaninate (**9**, 0.14 g) was dissolved in tetrahydrofuran (3 mL). The solution was added 37% aqueous hydrochloric acid solution (0.6 mL) slowly at 0 ºC, and the reaction mixture was allowed to warm to room temperature, and stirred for 2.5 h. After that, diluted with water (3 mL) and adjusted to pH = 8 by the addition of saturated aqueous sodium bicarbonate solution (6 mL), and extracted with ethyl acetate (3 mL). The organic layer was washed with brine (5 mL), and dried over anhydrous sodium sulfate, and evaporated. The obtained concentrate was subjected to silica gel chromatography eluting with solution containing 80% ethyl acetate and 20% methanol to afford the product **10** (0.07 g, 50%). $^1$H NMR (400 MHz, methanol-*d*$_4$) δ 8.24 (s, 1H), 8.20 (d, *J* = 15.2 Hz, 1H), 7.35 – 7.14 (m, 5H), 6.02 (d, *J* = 4.9 Hz, 1H), 4.62 (t, *J* = 5.1 Hz, 1H), 4.43 – 4.37 (m, 2H), 4.35 – 4.29 (m, 1H), 4.24 (br, 1H), 4.01 (dd, *J* = 10.9, 5.8 Hz, 1H), 3.96 – 3.90 (m, 2H), 1.45 (p, *J* = 6.3 Hz, 1H), 1.34 – 1.29 (m, 7H), 0.85 (t, *J* = 7.5 Hz, 6H) ; $^{13}$C NMR (101 MHz, methanol-*d*$_4$) δ 175.04, 174.98, 157.51, 157.45, 157.30, 157.25, 154.01, 153.99, 152.22, 152.15, 150.76, 150.74, 141.08, 130.80, 126.21, 126.19, 121.44, 121.39, 120.57, 90.09, 84.44, 84.36, 75.43, 71.66, 68.22, 67.67, 67.62, 51.65, 51.64, 49.88, 49.76, 49.69, 49.60, 49.55, 49.48, 49.39, 49.34, 49.26, 49.18, 49.12, 49.05, 48.91, 48.84, 48.63, 48.41, 41.78, 24.29, 24.27, 20.67, 20.61, 11.36, 11.32. FAB-MS (m/z): [M]+ calcd for $C_{25}H_{35}N_4{}^{15}N_2O_8P$, 580.6; found, 581.2.
